# Three-dimensional structure of a flavivirus dumbbell RNA reveals molecular details of an RNA regulator of replication

**DOI:** 10.1101/2020.09.17.300806

**Authors:** Benjamin M. Akiyama, Monica E. Graham, Zoe O′Donoghue, J. David Beckham, Jeffrey S. Kieft

## Abstract

Mosquito-borne flaviviruses (MBFVs) including dengue, West Nile, yellow fever, and Zika viruses have an RNA genome encoding one open reading frame flanked by 5′ and 3′ untranslated regions (UTRs). The 3′ UTRs of MBFVs contain regions of high sequence conservation in structured RNA elements known as dumbbells (DBs) that regulate translation and replication of the viral RNA genome, functions proposed to depend on the formation of an RNA pseudoknot. To understand how DB structure provides this function, we used x-ray crystallography and structural modeling to reveal the details of its three-dimensional fold. The structure confirmed the predicted pseudoknot and molecular modeling revealed how conserved sequences form a four-way junction that appears to stabilize the pseudoknot. Single-molecule FRET suggests that the DB pseudoknot is a stable element that can regulate the switch between translation and replication during the viral lifecycle by modulating long-range RNA conformational changes.

## INTRODUCTION

Mosquito-borne flaviviruses (MBFVs) are a widespread group of medically and economically important single-stranded positive-sense RNA viruses. Examples include dengue virus (DENV), the world′s most prevalent arbovirus that is endemic to over 100 countries and infects ∼390 million people annually (Bhatt et al., 2013), yellow fever virus (YFV), West Nile virus (WNV) and Zika virus (ZIKV) whose recent spread has demonstrated how rapidly MBFV outbreaks can emerge (Mackenzie et al., 2004). MBFV RNA genomes comprise a single open reading frame flanked by 5′ and 3′ untranslated regions (UTRs), important for regulating viral processes. The genomic RNA is both translated to yield viral proteins and is the template for genome replication, processes which cannot occur simultaneously on a single RNA as ribosomes moving in the 5′ to 3′ direction would collide with the replicase machinery moving in the opposite direction. To coordinate replication, these viruses use signals within the viral UTRs which can be identified by sequence and structure conservation and through mutagenesis studies demonstrating a replication defect when they are disrupted. Two such sequences in MBFV 3′ UTRs are conserved sequence 2 (CS2) and repeat conserved sequence 2 (RCS2) (Hahn et al., 1987). CS2 and RCS2 are contained within secondary structural elements called dumbbell (DB) RNAs that are proposed to contain tertiary structures called pseudoknots (Olsthoorn and Bol, 2001; Proutski et al., 1997; Proutski et al., 1999; Shurtleff et al., 2001) (Figure 1 – figure supplement 1). In dengue virus (DENV), two tandem DBs contain the RCS2 and CS2 motifs, respectively; however some flaviviruses have only a single DB (Villordo et al., 2016) (Figure 1A).

**Figure 1.**
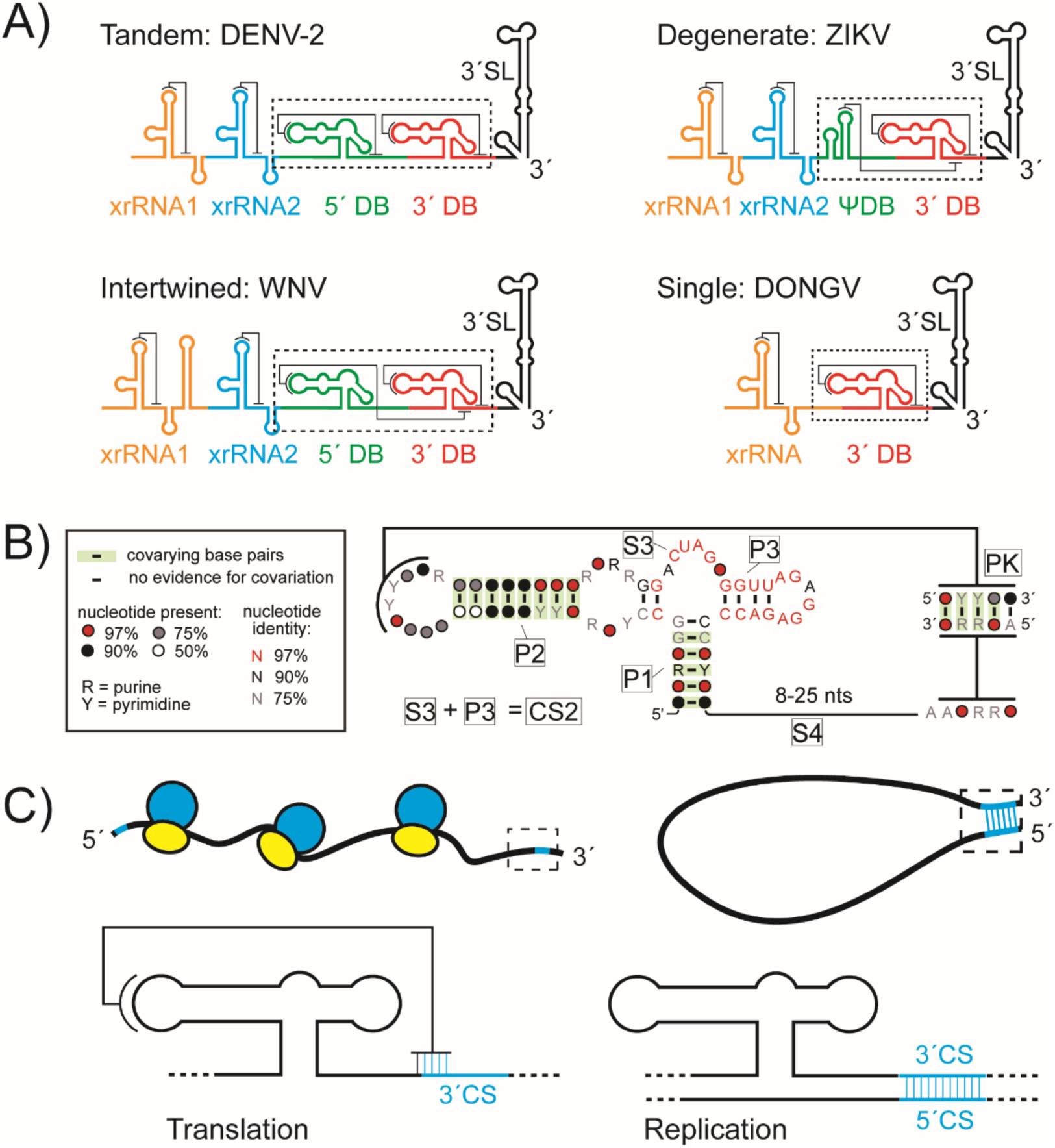
Conservation and role of flaviviral DB RNAs. **A)** Predicted organization of 3′ UTRs from DENV-2, West Nile virus, ZIKV, and DONGV. The 3′ dumbbell element is conserved but can exist in several configurations. **B)** Co-variation analysis of the 3′ DB element from 46 different viruses analyzed using R-scape (Rivas et al., 2017). Predicted helices (P1, P2, P3, and PK) are labeled as are predicted single-stranded linker regions (S3, S4). The S3 and P3 regions collectively make up a previously identified conserved sequence (CS2) (Hahn et al 1987). **C)** Model of the 3′ DB element during translation and replication. The pseudoknot overlaps with the 3′ CS, an element responsible for base pairing with the 5′ CS during viral minus-strand synthesis. During viral protein translation (left) the DB pseudoknot is formed. The pseudoknot is incompatible with 5′-3′ CS formation (cyan) during replication (right).

The importance of the DB elements, both virologically and clinically, has been established by a variety of studies. Functional investigations using mutagenized DENV replicons showed that DBs have a role in both viral protein production and replication, with particularly strong effects on regulation of replication (Alvarez et al., 2005a). Mutations that specifically disrupt pseudoknot base pairing demonstrated that this pairing was important for both translation and replication (Manzano et al., 2011). In DENV, the two DBs have distinct roles in viral replication in human and mosquito cells, suggesting DBs participate in a complex regulatory network of RNA-RNA interactions (de Borba et al., 2019). A 30 nucleotide deletion from the Dengue virus 4 DB (DENVΔ30) is highly attenuating (Durbin et al., 2001; Men et al., 1996), and has been developed into a vaccine candidate against all four DENV strains (Whitehead, 2016). Given that DB mutations are attenuating in all four DENV strains, a mechanistic understanding of the DB is of interest to public health efforts where strain-to-strain variation has hampered DENV vaccine development (Guzman et al., 2013). Understanding how a short truncation of a noncoding portion of the DB has such a large effect on pathogenicity could guide efforts to use diverse DB mutations for vaccine development (Shan et al., 2017).

Many RNA viruses regulate the switch between translation and replication states through changes in RNA structure. In the case of MBFVs, evidence suggests this switch involves a global conformational change in the genomic RNA, with translation occurring on a ‘linear’ genome conformation and replication on a ‘cyclized’ form with base pairing between the 5′ and 3′ ends (Alvarez et al., 2005b). Evidence for this includes conserved regions of co-varying base pairs between the 5′ and 3′ ends of the genome, known as the 5′ and 3′ conserved sequences (5′ and 3′ CSs) (Corver et al., 2003; Hahn et al., 1987; Khromykh et al., 2001; Lo et al., 2003). Disrupting the base pairing between the 5′ and 3′ CS severely affects viral replication, and atomic force microscopy showed cyclized RNAs with the viral replicase localized to the junction between the 5′ and 3′ ends (Alvarez et al., 2005b). Other interactions between the 5′ and 3′ ends supplement this base pairing, including the 5′ and 3′ ‘upstream of AUG regions’ and ‘downstream of AUG regions’ (Figure 1 - figure supplement 1) (Alvarez et al., 2008; Alvarez et al., 2005b; de Borba et al., 2015; Friebe et al., 2011; Liu et al., 2013). In addition, the location of a replication promoter at the genome′s 5′ end suggests that genome cyclization delivers the viral polymerase from its binding site at the 5′ end to signals at the 3′ end to begin replication (Davis et al., 2007; Filomatori et al., 2006; You et al., 2001).

The location of the DBs and their relationship to other elements in the 3′ UTR suggests these structures play important regulatory roles. Specifically, the DBs are directly adjacent to the 3′ CS and in the majority of MBFVs the pseudoknot directly overlaps with the 3′ CS. This led to the hypothesis that the DB pseudoknot could act as a sensor of 5′-3′ CS base pairing, or act as a competitor to cyclization, temporally repressing viral replication to maximize viral protein translation (de Borba et al., 2019). In support of this assertion, a simplified DENV minigenome containing only the 5′ and 3′ UTRs demonstrated that the most downstream DB (3′ DB) RNA pseudoknot does not form when the RNA is cyclized (Sztuba-Solinska et al., 2013). This idea is further supported by an analysis of sequence variants in MBFVs and in revertant strains of CS mutants, showing that the 3′ DB pseudoknot regulates genome cyclization during replication (de Borba et al., 2019). Finally, it was proposed that the most upstream DB (5′ DB) from DENV competes with an alternate conformation that base pairs with a region of the capsid protein coding region during viral replication (Figure 1 – figure supplement 2) (de Borba et al., 2015). Collectively, this suggests a model in which the DB pseudoknots compete with the cyclized form of the genome. DBs have also been implicated in the formation of non-coding subgenomic flaviviral RNAs (sfRNAs), which are formed through resistance to degradation by the host 5′→3′ exonuclease XRN1, using RNA structures in the 3′ UTR known as XRN1-resistant RNAs (xrRNAs) (Chapman et al., 2014a). It has been theorized that DBs, by virtue of their complex secondary structures and tertiary interactions, may act as xrRNAs, but this requires further testing.

The presence of conserved secondary and tertiary structural features in MBFV DB elements suggests that their three-dimensional fold dictates function (Villordo et al., 2016), but until now this structure was unknown. We therefore used x-ray crystallography to determine the three-dimensional structure of a DB from Donggang virus (DONGV), an insect-specific flavivirus with homology to many medically important MBFVs. The RNA crystallized as a dimer by a domain exchange between CS2 elements, but both copies had folded RNA pseudoknots and the structure provided sufficient information to model the monomeric form. This model contains a four-way junction in which the CS2 sequence folds into two helices, imparting stability to the overall fold and forming an available surface for potential protein interactions. Chemical probing of the DB structure supports this model and shows the same structure is formed by DENV DBs. Using smFRET experiments to explore conformational dynamics, we discovered that the DB can act as a switch and that the pseudoknotted form is heavily favored under physiologic conditions, with implications for its function as a regulator of viral replication.

## RESULTS

### Base pairing covariation identifies a conserved dumbbell RNA in the viral 3′ UTR

We first bioinformatically analyzed MBFV 3′ UTRs to identify conserved secondary structures and features of DBs from diverse sources. Overall, the 5′ DB is more variable between species and can be absent or altered compared to the 3′ DB (Figure 1A). We therefore hypothesized that the 3′ DB is more important, an idea supported by the presence of CS2 within it (Hahn et al., 1987). To assemble a list of homologous DB structures, we started with a small sequence alignment of 3′ DB RNAs, identified 46 putative DB sequences by RNA base pairing covariance using the Infernal algorithm (Nawrocki and Eddy, 2013), then aligned them using R-scape (Rivas et al., 2017). This identified helices (P1 and P2) by base pair covariance (Figure 1B). A third helix (P3) was predicted to fold by the mFold and Vienna RNA secondary structure prediction algorithms (Gruber et al., 2008; Zuker, 2003), however its sequence is so conserved that there is no base pairing covariation. Manual alignment of the sequences identified co-varying base pairs consistent with an RNA pseudoknot in all the DB sequences (Figure 1B). This pseudoknot often overlaps with the 3′ CS; in such cases the pseudoknot cannot form when the 5′ and 3′ CSs base pair during viral replication (Figure 1C) (de Borba et al., 2019).

### The DB crystal structure reveals a complex three-dimensional fold and RNA pseudoknot

To understand how DB structure may participate in global RNA conformational changes, we used x-ray crystallography to reveal its three-dimensional structure. After screening DBs from multiple MBFVs, we obtained diffracting crystals using a sequence from DONGV, an insect-specific flavivirus (Blitvich and Firth, 2015; Takhampunya et al., 2014). The DONGV DB sequence is conserved with other MBFVs including DENV, WNV, YFV, and ZIKV, thus structural information from DONGV DB is broadly applicable (Figure 1 – figure supplement 3). DONGV DB RNA crystals were soaked with iridium (III) hexammine, which was used to determine phases and solve the three-dimensional structure (Keel et al., 2007). The final structural model was refined to a resolution of 2.1 Å, providing high-resolution information (Table 1).

**Table 1.** Crystallographic data collection and refinement statistics

|  |  |
| --- | --- |
| **Data collection** |  |
| Space group | P2<sub>1</sub> |
| Cell dimensions |  |
| *a*, *b*, *c* (Å) | 65.93, 49.67, 73.46 |
| $\alpha$, $\beta$, $\gamma$ (°) | 90, 99.73, 90 |
| Wavelength (Å) | 1.102 |
| Resolution (Å) | 40.96 - 2.1 (2.175 - 2.1) |
| *R*<sub>merge</sub> | 0.1303 (0.8773) |
| R-meas | 0.1367 (0.9256) |
| R-pim | 0.04105 (0.2914) |
| *I*/σ*I* | 17.95 (3.17) |
| Completeness (%) | 99.81 (100.00) |
| Multiplicity | 10.9 (10.0) |
| CC1/2 (%) | 0.998 (0.843) |
| CC* | 0.999 (0.956) |
| Wilson B-factor | 34.69 |
| **Refinement** |  |
| Resolution (Å) | 40.96 - 2.1 (2.175 - 2.1) |
| No. of unique reflections | 27599 (2744) |
| *R*<sub>work</sub> / *R*<sub>free</sub> | 0.21/0.26 |
| CC<sub>work</sub>/CC<sub>free</sub> | 0.95 (0.81) / 0.93 (0.74) |
| No. atoms |  |
| RNA | 3822 |
| Ligand/ion | 358 |
| Water | 89 |
| B-factors |  |
| RNA | 33.39 |
| Ligand/ion | 45.38 |
| Water | 28.92 |
| R.m.s deviations |  |
| Bond lengths (Å) | 0.038 |
| Bond angles (°) | 1.53 |
| Clashscore | 4.91 |
| Coordinate error | 0.33 |
| Phase error | 29.78 |

\*Highest resolution shell is shown in parenthesis.

The DONGV DB crystallized as a dimer in the asymmetric unit (Figure 2A and Figure 2 - figure supplement 1A), with both copies adopting a similar fold (Figure 2 - figure supplement 1B). As predicted, helices P1, P2, and the pseudoknot formed in both copies. However, the predicted P3 stem did not form, instead its sequence made base pairs with a linker predicted to be single stranded (S3) (Figure 2A and 2B).

**Figure 2.**
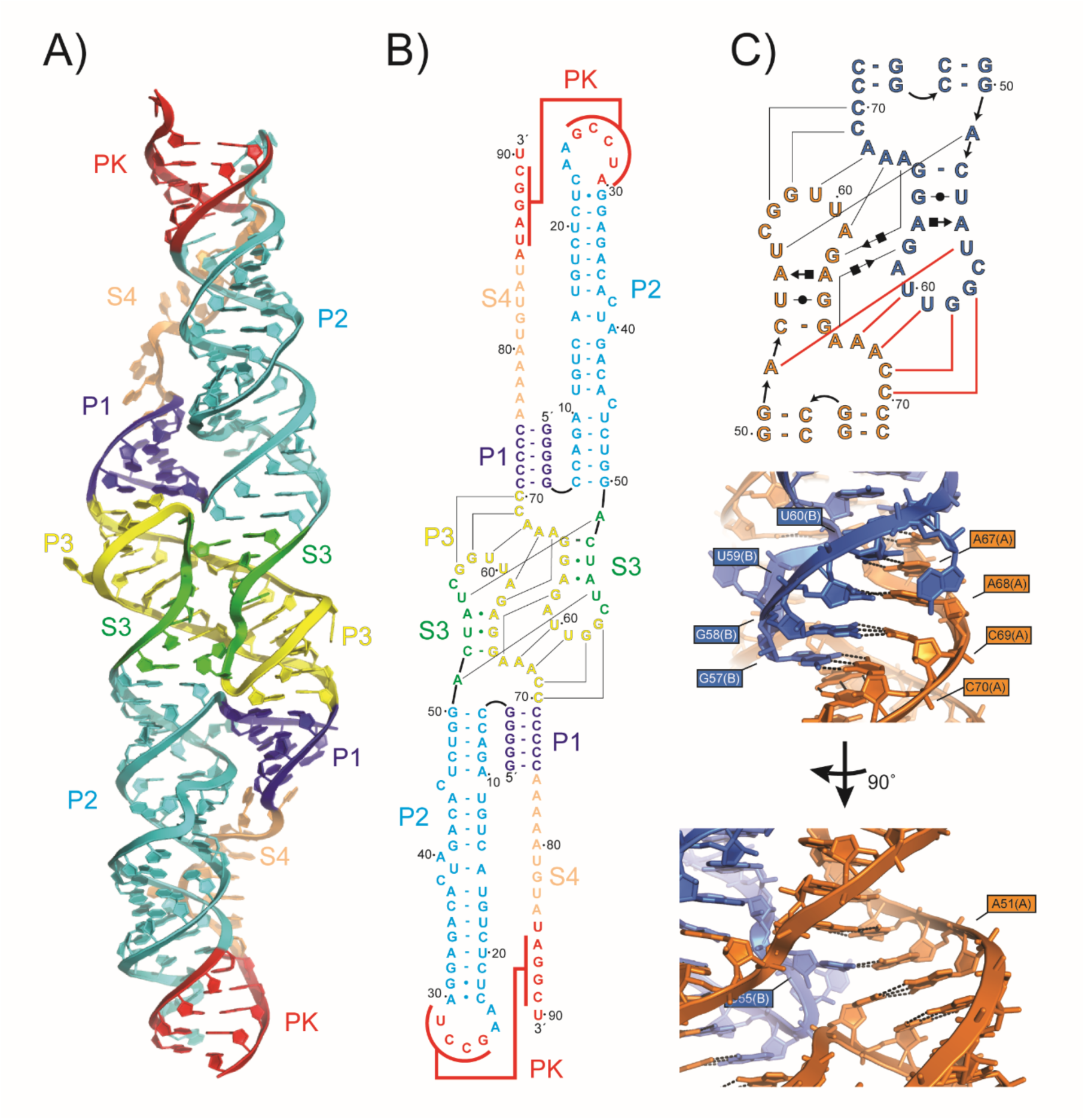
Crystal structure of the DONGV DB RNA. **A)** Overall structure of the dimer form of the DONGV DB observed in the crystal. Iridium (III) hexammine ions used for SAD phasing are omitted for clarity. Predicted stems from Figure 1B are labeled (P1 = blue, P2 = cyan, P3 = yellow, pseudoknot = red) as are the predicted single-stranded regions (S3 = green, S4 = orange). **B)** Base pairing interactions observed in the RNA crystal structure colored as in (A). Lines indicate base pairing observed between copy 1 and copy 2. **C)** Secondary structure in Leontis-Westhof notation (top) and detailed three-dimensional view (bottom) of the dimer interface highlighting *in trans* base pairing interactions between copy 1 (orange) and copy 2 (blue). Red lines in the secondary structure show base pairing interactions labeled in the three-dimensional view.

Additional intermolecular base pairs also formed in this region, forming an extensive interface of crystal contacts (Figure 2B and C). The largest difference between the two molecules in the crystallographic dimer was in the single-stranded RNA linker S4 and there was a slight difference in the orientation of the P2 helix (Figure 2 – figure supplement 1B).

Both copies of the DONGV DB observed in the asymmetric unit formed the predicted pseudoknot with at least four base pairs, but the two copies differed somewhat in this region (Figure 3A and B). Copy 1 (chain A in the deposited coordinate file) formed four Watson-Crick (WC) base pairs in the pseudoknot, however the base pair predicted to form between A30 and U85 was not observed (Figure 3A). Instead, A30 participates in an A•C interaction with C23, which is positioned for this interaction by adjacent residues in the P2 loop. A24 stacks underneath C23, participating in an A-U-A base triple interaction with the U29-A86 base pair. U85 stacks on A84, which hydrogen bonds to the 2′OH of G31 (Figure 3B). In contrast, copy 2 (chain B in the deposited coordinate file) has five base pairs in its pseudoknot with the A30-U85 base pair formed as predicted, and copy 2 has neither the A•C base pair nor the A-U-A base triple observed in copy 1 (Figure 3C). The distinct pseudoknot organizations in copies 1 and 2 may represent two alternate conformations of the RNA captured in the crystal.

**Figure 3.**
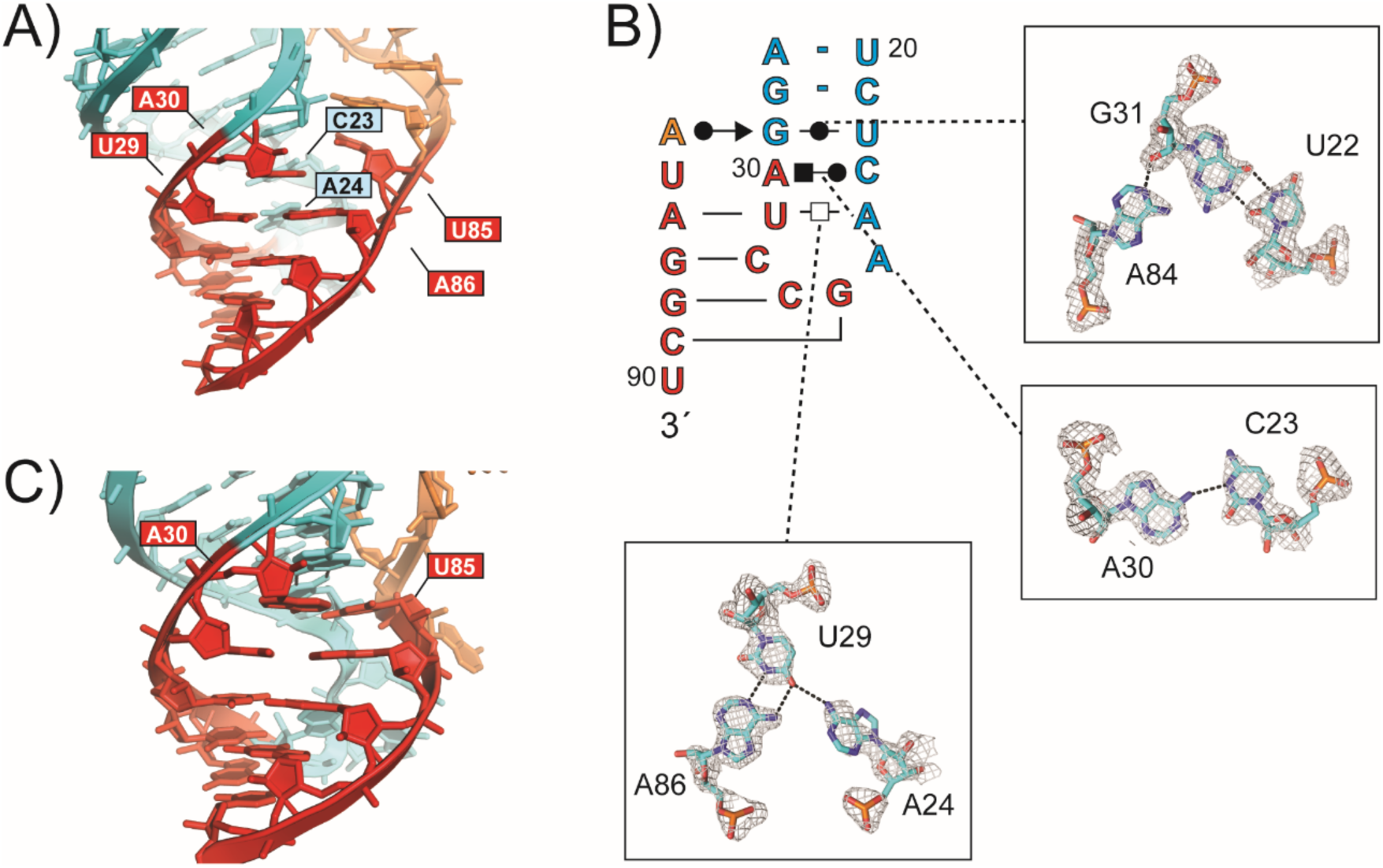
Molecular details of the RNA pseudoknot. **A)** Detailed view of the DB RNA pseudoknot in copy 1 of the asymmetric unit colored as in Figure 2A. Nucleotides A30 and U85 are labeled, this predicted base pair is not formed. The position of A30 is stabilized by non-canonical base pairing interactions in the P2 stem-loop (Figure 3B). The position of iridium (III) hexammine ions used for phasing are shown as magenta spheres, with the amines not visualized for clarity. **B)** Secondary structure in Leontis-Westhof notation of copy 1 pseudoknot loop, insets show details of non-canonical base pairing stabilizing the loop in the absence of the A30-U85 base pair with electron density from a composite omit map of the crystal shown at the 1σ contour level. **C)** Detailed view of the pseudoknot of copy 2. A30 and U85 are labeled and are base paired in this copy.

### Molecular modeling of a monomeric structure

Examination of each copy of the DB individually, outside of the context of the crystal, revealed distorted base pairing interactions in the S3 and P3 regions with far from ideal A-form RNA geometries and large internal solvent channels (Figure 2 – figure supplement 2B). This suggests that crystal packing in this region distorts the RNA fold compared to its solution conformation. Close examination of the intermolecular crystal contacts revealed that the predicted base pairs in the P3 stem formed, but *in trans*, as a domain-swap between the two copies of the RNA (Figure 2C). Domain-swapping is frequently observed in crystallography of both proteins and RNA, due to the high concentration of molecules when naturally occurring interactions that occur *in cis* at physiologic concentrations form *in trans* in the crystal (Chapman et al., 2014a; Liu and Eisenberg, 2002; Pfingsten et al., 2006; Steckelberg et al., 2018). We therefore hypothesized that the P3 stem is formed in solution *in cis,* but forms *in trans* in the crystal due to crystal contacts. We used the fact that the base pairing arrangement of the consensus model was observed in the crystal, albeit in a domain-swapped interaction, to model the monomer structure as it would occur *in cis.* We excised nucleotides 55-63 from copy 2 and transferred them to copy 1, and *vice versa,* to model the P3 stem formed *in cis* (Figure 4 – figure supplement 1). The remainder of the RNA structure is consistent with this arrangement and few other perturbations were required to create the model, other than repositioning the connecting nucleotides 54 and 64. These two were deleted and positioned without using electron density to connect the two chains, producing a monomeric model of the RNA guided by the crystal structure (Figure 4).

**Figure 4.**
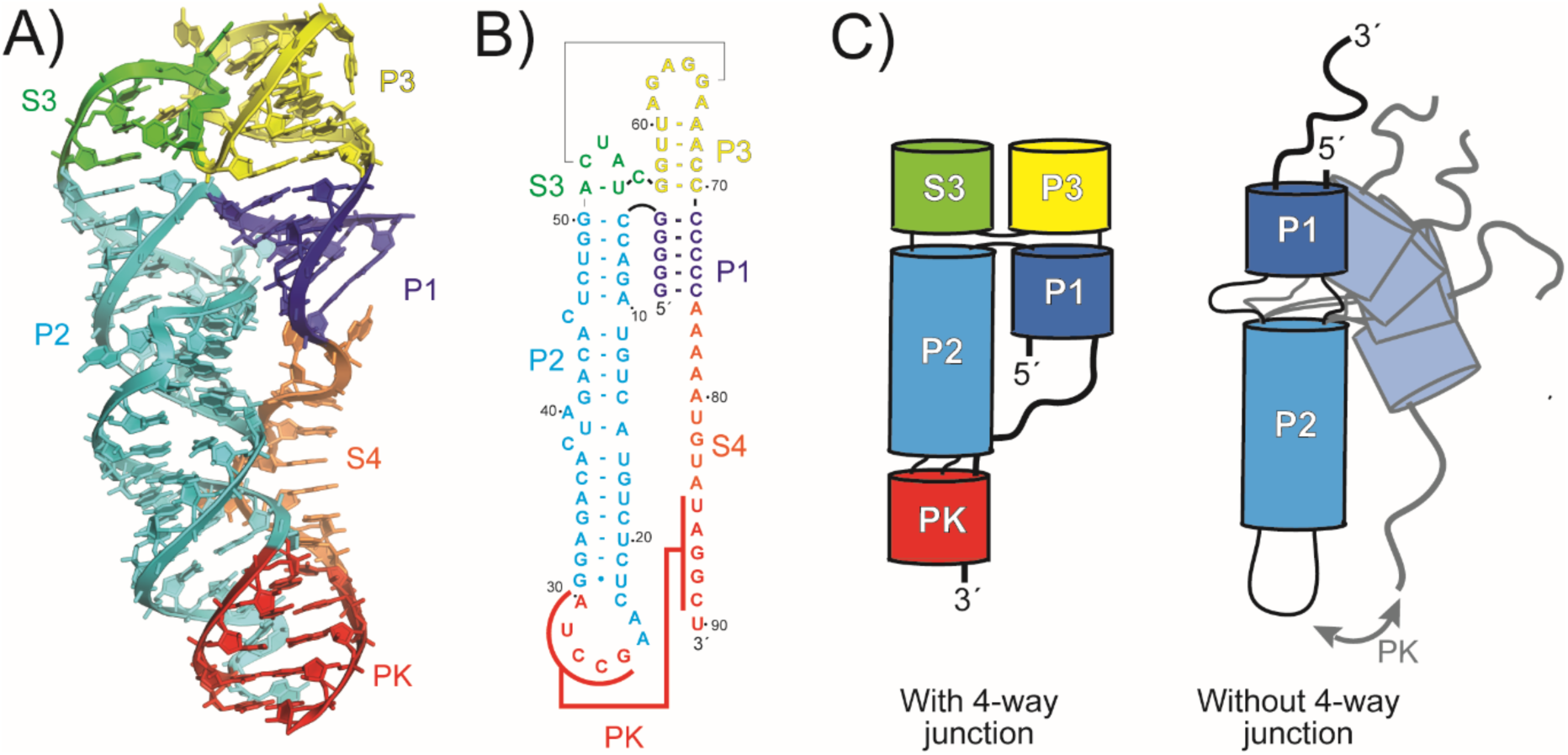
Model of the monomeric DB RNA. **A)** Three-dimensional model of the monomeric solution structure of the DONGV DB guided by the crystal structure. The structure forms a four-way junction with co-axial stacking of the P1 (blue) and P3 (yellow) stems and with the P2 (cyan) and S3 (green) stems. **B)** Secondary structure map of the DONGV DB solution structure model. **C)** Cartoon model demonstrating a four-way junction with limited conformational space (left) compared against an RNA without a four-way junction (right) that can sample many different conformations.

### Chemical probing supports the monomeric structure model of the DB RNA

We used chemical probing to test if the resultant modification patterns are consistent with our monomeric structural model. Wild-type DONGV and DENV-2 DBs were used as well as a DONGV DB with a pseudoknot disrupted by mutation. We used N-methylisatoic anhydride (NMIA), which detects flexible regions of the RNA backbone, as a proxy for unpaired nucleotides (Merino et al., 2005), and dimethyl sulfate (DMS), which detects N1 on adenines and N3 on cytosines that are not hydrogen bonded (Inoue and Cech, 1985). The reactivity pattern was consistent with the monomeric structural model in the P3 and S3 regions (Figure 5A and Figure 5 – figure supplement 1, Figure 5 – figure supplement 2, Figure 5 – figure supplement 3), with high reactivity in loop regions and low reactivity in Watson-Crick base-paired regions. In contrast, the reactivity pattern did not agree with the dimeric form observed in the crystal in the P3 and S3 regions, providing strong evidence that the monomeric structural model more accurately represents the folded solution state. Probing of the DONGV DB pseudoknot mutants confirmed the formation of the pseudoknot (Figure 5C). Specifically, residues 26-30 were mutated to their complementary sequence (PK_mut1_) as well as residues 85-89 (PK_mut2_), and a third RNA contained both mutations to restore the pseudoknot by forming compensatory base pairs (PK_comp_). As predicted PK_mut1_ and PK_mut2_ showed higher reactivity in the regions forming the pseudoknot, consistent with pseudoknot disruption, while the PK_comp_ mutant′s reactivity pattern resembled wild-type RNA.

**Figure 5.**
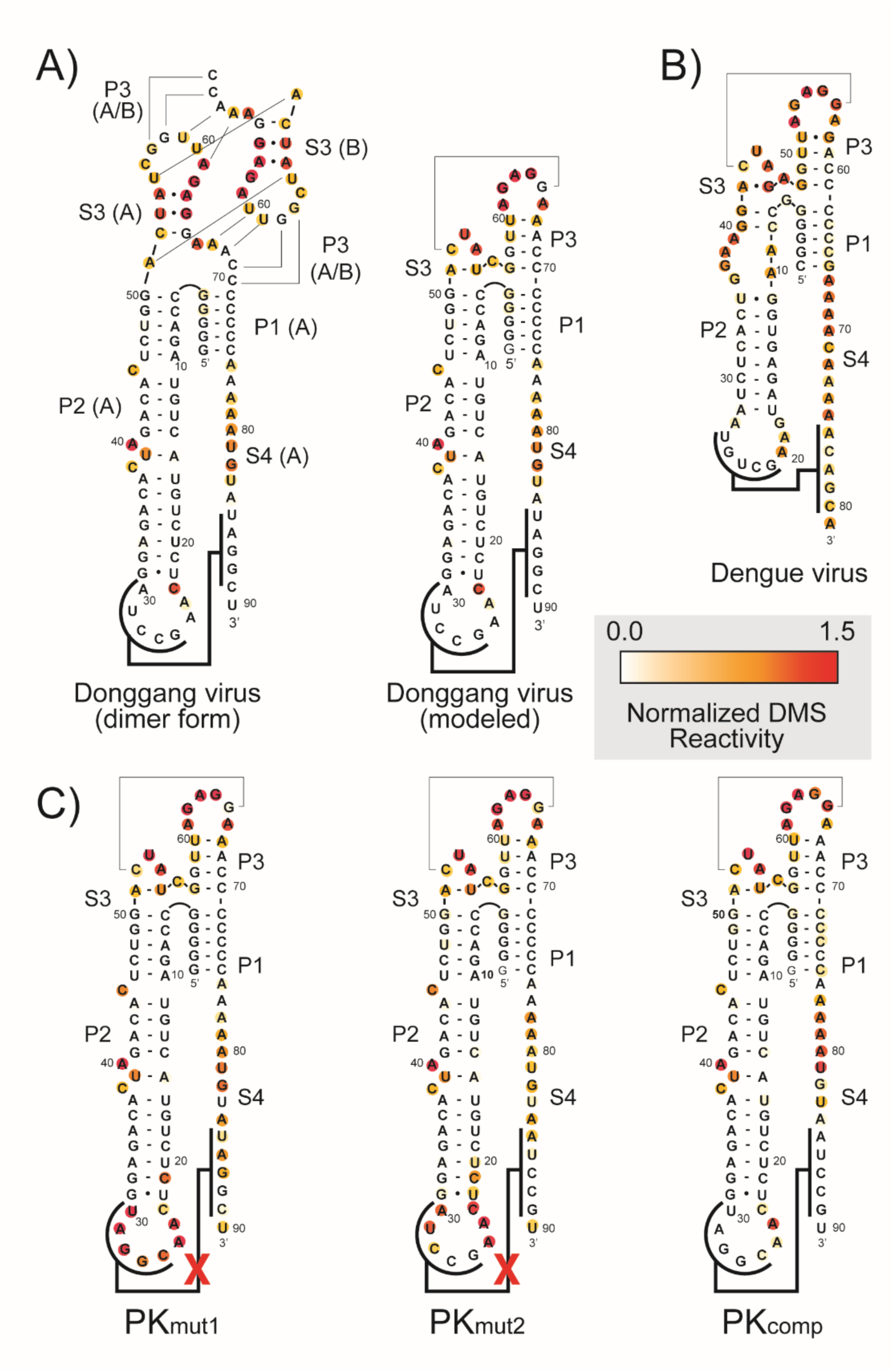
NMIA chemical probing of the DB RNA. **A)** Secondary structure of the DONGV DB RNA in the form observed in the crystal structure (dimer form) compared with the modeled form. NMIA reactivity of the wild-type DB sequence is color-coded (red is high reactivity, white is low reactivity). **B)** NMIA reactivity of the DENV-2 DB plotted on the putative secondary structure based on homology to DONGV. **C)** NMIA reactivity of pseudoknot mutant RNAs (PK_mut1_ and PK_mut2_) plotted on the modeled RNA secondary structure. The mutations in PK_mut1_ and PK_mut2_ were combined to generate compensatory base pairing (PK_comp_).

We next probed a DB from DENV-2 to determine if the RNA fold is conserved. The similar reactivity profiles between DONGV and DENV-2 indicated that the overall folding of the RNA is maintained (Figure 5A and B and Figure 5 – figure supplement 1). In particular, both had reactivity patterns consistent with the formation of the P1, P2, and P3 stems and were protected in their pseudoknots, suggesting that the pseudoknot was formed. This, combined with the conservation in the consensus model (Figure 1B), strongly supports that the DONGV DB structure solved here is representative of DB elements across the flaviviruses.

### The modeled monomeric RNA structure contains a four-way junction

A defining feature of the DONGV DB RNA structure is an acute angle between the P1 and P2 stems, which appears to be important for favoring pseudoknot formation. This angle is associated with a sharp kink in the RNA backbone that places the 3′ single-stranded region (S4) in close proximity to the P2 stem-loop (Figure 4A and B). In the crystal structure, several bases in the S3 and P3 regions stack on the P1 and P2 helices. As base stacking is central to stabilizing folded RNA structures, these interactions likely stabilize the global architecture of the DB, including the acute angle between P1 and P2. In the monomeric structure model, the stacking pattern of P1 and P2 with S3 and P3 create a four-way junction (Figure 2C). Specifically, in S3 the A51-U55 pair stacks on the P2 stem, and the C52-G65 pair stacks on A51-U55. The C52-G65 base pair may help lock the angle between the S3 and P3 stems (Figure 6A and B). The P3 stem in the model stacks on the P1 stem, completing the four-way junction. Comparing the model solution structure with existing RNA structures reveals that the four-way junction can be classified as a member of family H (Laing and Schlick, 2009) with homology to a four-way junction in RNase P that stabilizes the specificity domain (Figure 6 – figure supplement 1) (Krasilnikov et al., 2003). Overall, the model suggests a hypothesis for the stability of the DB pseudoknot: the four-way junction structure locally favors the acute angle between P1 and P2, which globally brings the two complementary sequences of the pseudoknot in proximity (Figure 4C).

**Figure 6.**
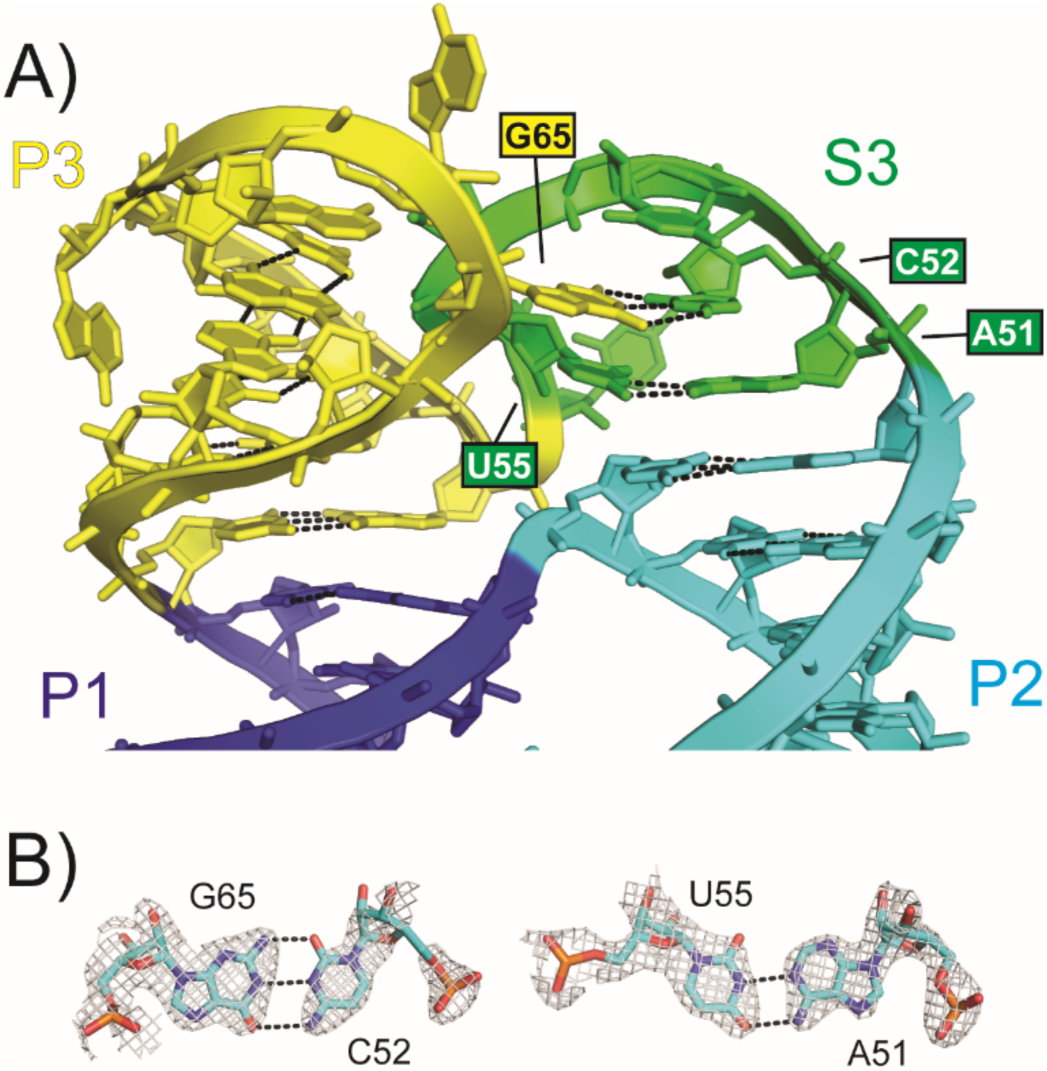
Molecular details of the modeled four-way junction. **A)** Detailed view of the four-way junction from the modeled monomeric RNA. The four-way junction is formed by base stacking between the P1 (blue) and P3 (yellow) stems and between the P2 stem (cyan) and S3 (green) stems. Base pairs between A51 and U55 and between C52 and G65 are labeled. **B)** Molecular details of the base pairs highlighted in (A). Electron density is from a composite omit map of the crystal structure at 1σ contour level.

### The DB pseudoknot is the heavily favored conformation

The observation that both DB copies in the crystal formed the pseudoknot combined with chemical probing indicating a pseudoknot in solution suggest that this tertiary interaction is favored, an idea further supported by the presence of pervasive base stacking and base pairing throughout the structure and a four-way junction that may stabilize a conformation that facilitates pseudoknot formation. To test this idea, we used single-molecule Förster resonance energy transfer (smFRET) to observe pseudoknot dynamics with temporal resolution on individual molecules. A Cy3 fluorophore was attached to the DONGV DB in the P2 stem (Figure 7A and Figure 7 – figure supplement 1A), and a 3′ extension was added that is complementary to a DNA oligonucleotide used to immobilize the complex on a microscope coverslip using a biotin-streptavidin linkage (Figure 7A and Figure 7 – figure supplement 1A). This DNA oligo was modified with Cy5 at a position designed to display higher FRET when the pseudoknot is formed relative to that when it is not formed. The Cy3-modified RNA was annealed to the Cy5-modified DNA oligo and immobilized on a microscope coverslip followed by smFRET observation of real-time RNA folding (Figure 7A). The vast majority of individual spots that photobleached had single photobleaching events, suggesting that each spot on the microscope slide had only a single Cy3 and Cy5 molecule, further demonstrating that the DB is monomeric in solution (Figure 7 – figure supplement 2).

**Figure 7.**
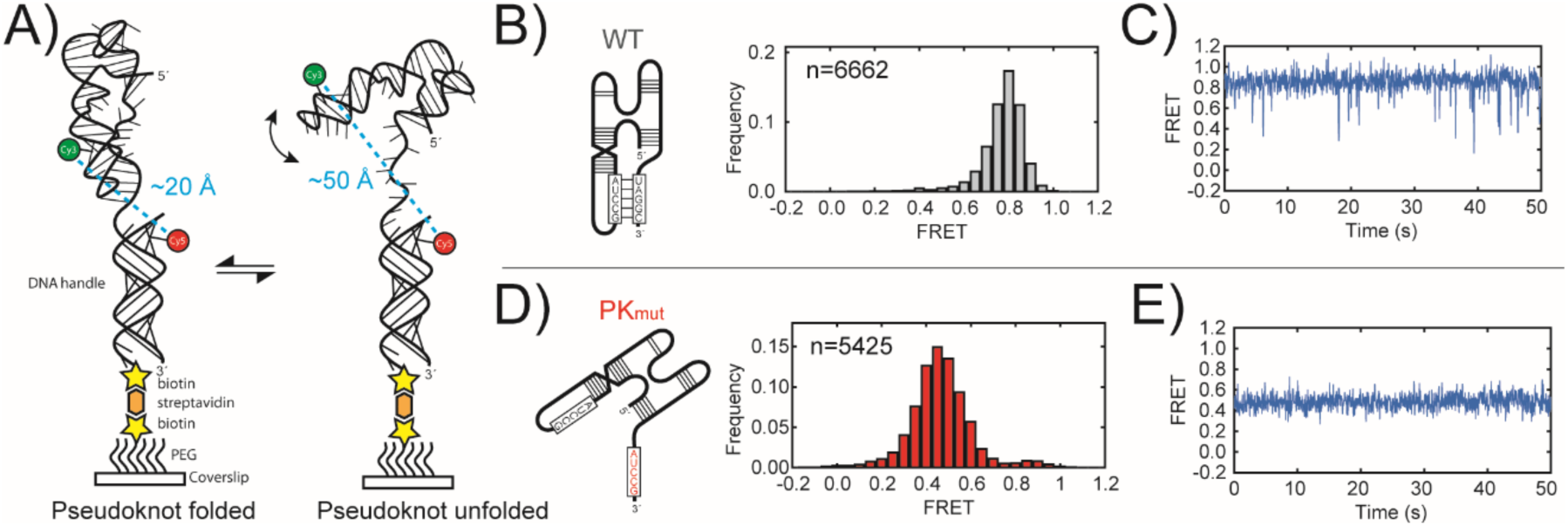
smFRET measurements of pseudoknot stability. **A)** Labeling strategy for smFRET experiments. DONGV DB RNA was site-specifically labeled with Cy3. A 3′ extension on the RNA was used to immobilize the RNA construct on a microscope slide based on hybridization to a biotinylated and Cy5-labeled DNA handle. A folded pseudoknot is predicted to have a high FRET value and an unfolded pseudoknot is predicted to have a lower FRET value. **B)** smFRET histogram of wild-type DB RNA. (left) Model of the RNA construct. (right) Histogram of the distribution of FRET values, n=number of molecules observed. **C)** Representative smFRET trace of a wild-type DB RNA. **D)** smFRET histogram of PK_mut_ DB RNA. (left) Model of the RNA construct used. (right) smFRET histogram. **E)** Representative smFRET trace of a PK_mut_ DB RNA.

smFRET histograms compiled from wildtype DONGV DBs showed a single Gaussian distribution centered at 0.8 FRET (Figure 7B). To test whether this FRET state corresponded to the pseudoknotted form, we generated a pseudoknot mutant (PK_mut_) RNA and tested it in this assay. The PK_mut_ RNA showed a single predominant FRET distribution at ∼0.5 FRET, verifying that this FRET state corresponds to the unfolded form of the pseudoknot and that the high FRET state observed in the wild-type RNA is the pseudoknotted form (Figure 7D). Thus, smFRET distributions from the wildtype RNA showed that the folded pseudoknot is the heavily favored conformation.

We next examined the conformational dynamics of individual DB molecules, revealed in the smFRET traces as a function of time. As expected, PK_mut_ single-molecule traces displayed a stable 0.5 FRET value without any detectable transitions to the higher FRET value (Figure 7E and Figure 7 – figure supplement 1C), consistent with the pseudoknotted state being largely destabilized by the mutations. In contrast, wild-type traces showed molecules predominantly in the folded 0.8 FRET state, with only occasional transient excursions to a 0.5 FRET state that are so short-lived as to be near the limit of detection of the camera (∼0.1 seconds) (Figure 7C and Figure 7 – figure supplement 1B). The short duration of the unfolded 0.5 FRET state showed that breaking the pseudoknot interaction is followed by rapid re-formation, consistent with a low kinetic barrier in the transition from the unfolded to folded state (Figure 7 – figure supplement 3). However, the distribution of states revealed that the pseudoknotted state predominates and therefore is thermodynamically much more stable than the unfolded state, with only a small population observed in the unfolded configuration. The transient excursions to a lower FRET state were apparent in conditions containing 2 mM MgCl_2_, however they were not apparent at 10 mM MgCl_2_, demonstrating Mg^2+^′s role in stabilizing the pseudoknot fold (Figure 7 – figure supplement 4). This is expected, as Mg^2+^ folding dependence is a common feature of RNA tertiary structure (Grilley et al., 2006).

The DB pseudoknot appears to be more stable than many other pseudoknots measured in biophysical assays including the human telomerase RNA pseudoknot (Hengesbach et al., 2012), the PreQ1 riboswitch pseudoknot (Warnasooriya et al., 2019), and the *dianthovirus* XRN-resistant RNA pseudoknot (Steckelberg et al., 2018). Since the DB pseudoknot is mutually exclusive with long-range 5′-3′ base pairing associated with viral replication (Figure 1C), the stability of the DB pseudoknot suggests that the DB folding landscape favors the translation-competent and not replication-competent form of the viral genome in the DB′s role as a sensor or regulatory element.

### Dumbbell RNAs do not efficiently resist the 5**′**→3**′** exonuclease Xrn1

xrRNA elements in the flaviviral 3′ UTR form three-dimensional folds that resist degradation by host 5′→3′ exonucleases, predominantly the host protein XRN1 (Funk et al., 2010; Pijlman et al., 2008). These xrRNAs form a structure where the 5′ end is surrounded by a distinctive ring-like fold, forming a “molecular brace” preventing the 5′ end from being pulled into the XRN1 active site (Akiyama et al., 2016a; Akiyama et al., 2016b; Chapman et al., 2014a), creating sfRNAs (Pijlman et al., 2008). It has been proposed that DBs may also be XRN1 resistant, as lower molecular weight sfRNAs that could correspond to XRN1 stalled at DB RNA elements have been observed (Funk et al., 2010). However, *in vitro* XRN1 resistance assays and northern blots of YFV infections suggest that these low molecular weight sfRNAs may be due to trimming of the 3′ end of the sfRNA, not DB exonuclease resistance. (Chapman et al., 2014b; Silva et al., 2010).

The DONGV DB 5′ end is surrounded by other RNA elements within the structure (Figure 4B), but lacks the distinctive topology observed in xrRNAs, in which a continuous strand forms a ring of defined size through which the 5′ end is threaded (Akiyama et al., 2016b). Therefore, we tested DB Xrn1 resistance using a ZIKV model system with an established assay in which *in vitro* transcribed RNA is modified to contain a 5’ monophosphate and a 29 nucleotide 5’ leader to allow recombinant purified K. lactis Xrn1 to load on the 5’ end to begin degradation (Chapman et al., 2014b). In 2 mM MgCl_2_, the ZIKV xrRNA was highly resistant to Xrn1, with a significant majority of the input RNA shifted to a lower molecular weight band corresponding to the partially degraded RNA. In contrast, at 2 mM MgCl_2_ the ZIKV DB was efficiently degraded and no Xrn1-resistant band was observed (Figure 8A). Interestingly, at 10 mM MgCl_2_ an Xrn1-resistant fragment was observed in the DB, albeit at a much lower intensity (∼20% of input). This is consistent with high magnesium concentrations stabilizing the DB fold (Figure 7 – figure supplement 4), although even at these higher magnesium concentrations only a small amount of Xrn1 resistance was observed. Therefore, while the DB elements are stable and complex structures containing a pseudoknot, we find that they do not efficiently resist Xrn1, consistent with the lack of a tight ring-like conformation through which the 5′ end is threaded.

**Figure 8.**
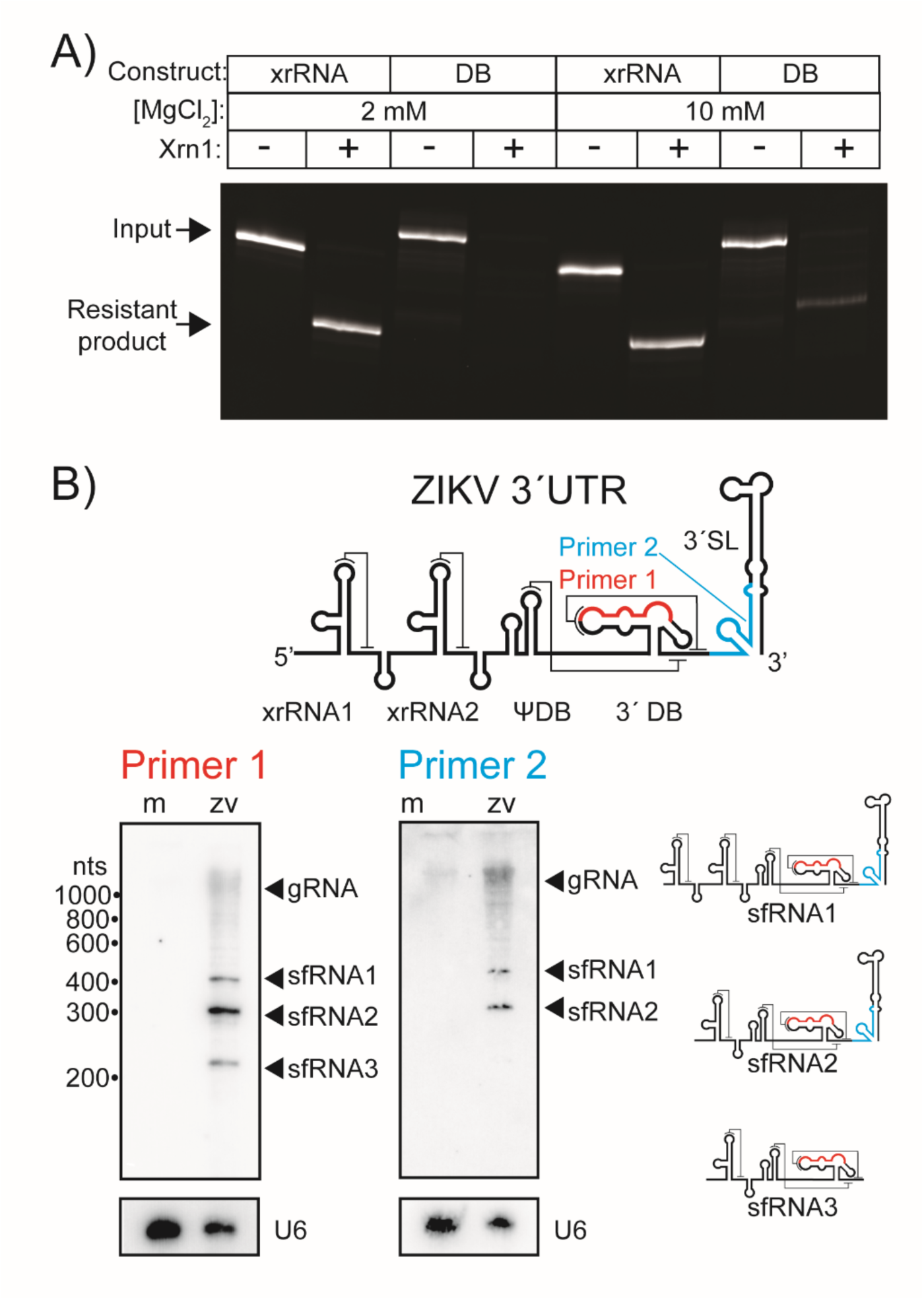
DB resistance to the 5′→3′ exonuclease Xrn1. **A)** Xrn1 digest of ZIKV Xrn1-resistant RNA (xrRNA) or ZIKV DB RNA in the presence of 2 mM or 10 mM MgCl_2_. The upper bands correspond to RNAs with an intact 5′ single-stranded leader that have not been treated with Xrn1. The lower bands correspond to RNA fragments resistant to Xrn1 digestion, in which Xrn1 has removed the 5′ single stranded linker leaving the resistant fragment behind. **B)** Upper panel: the 3′ UTR of ZIKV highlighting the position that northern blot DNA probes hybridize to for primer 1 (red) and primer 2 (cyan). Lower panel: northern blots from identical mock (m) and ZIKV infected (ZV) cellular RNA probed with either primer 1 or primer 2. Blots were also probed with a DNA oligo hybridizing with the U6 snRNA as a loading control. Positions of the genomic RNA (gRNA) and subgenomic flaviviral RNAs (sfRNAs) are as labeled. Cartoon diagrams of sfRNA1, sfRNA2, and sfRNA3 are shown.

Given their limited Xrn1 resistance in *in vitro* assays, we next determined if MBFV DBs demonstrate XRN1 resistance during infection, as infection with DENV, WNV, and ZIKV results in two sfRNAs that are clearly due to two xrRNAs, but other lower molecular weight bands as well (Akiyama et al., 2016b; Chapman et al., 2014a; Funk et al., 2010). These ′extra′ sfRNAs could be formed by one of two possible mechanisms: either XRN1 resistance of the DBs or trimming of the 3′ end by another nuclease. To distinguish between these mechanisms, we infected A549 cells with ZIKV and detected sfRNAs using northern analysis with two different oligonucleotide probes: one hybridized within the DB and a second hybridized within the terminal 3′ stem-loop. If the DBs are XRN1 resistant during infection, the same sfRNA fragments should be observed with both probes. Conversely, if additional sfRNA species are formed by trimming of the 3′ end, the northern blot with a probe directed against the 3′ stem-loop should have fewer bands than the northern blot probed against the DB. With the probe directed against the DB, three predominant sfRNAs were observed—sfRNA1, sfRNA2, and sfRNA3—with sfRNA1 and sfRNA2 formed due to XRN1 resistance of xrRNA1 and xrRNA2 (Figure 8B). With the probe against the 3′ stem-loop, only two sfRNAs were observed, sfRNA1 and sfRNA2 (Figure 8B). This clearly demonstrates that sfRNA3 is formed due to 3′ trimming of the viral sfRNA and not due to XRN1 resistance of the DB, consistent with the lack of a defined ring-like fold in the DBs and the *in vitro* assay.

## DISCUSSION

Here we describe the three-dimensional structure of the DONGV DB, an important functional RNA element in the flaviviral 3′ UTR that is implicated in regulating viral replication and whose structural disruption has been linked to attenuated viruses. Using our crystal structure, structure-guided molecular modeling, solution-based chemical probing and single-molecule experiments, we determined that the DB contains a four-way junction, with a global architecture that favors the formation of an unusually stable pseudoknot associated with regulating replication.

Within the structure, numerous interactions in addition to pseudoknot base pairing appear to favor a global conformation that promotes pseudoknot formation. That is, the overall DB conformation is such that it brings the sequences that pair to form the pseudoknot into proximity. The features that favor this fold likely include a four-way junction, which appears to constrain two of the emergent helices into an acute angle.

Indeed, helical junctions are critical determinants of global RNA structure as they dictate the angle between emerging helices through the formation of a specific pattern of base stacking (Laing and Schlick, 2009; Lescoute and Westhof, 2006). By favoring specific angles, junction-induced global organization brings distal regions near each other in space such that long-range tertiary interactions can form. This structure suggests this principle of RNA folding operates in the DBs.

Our discovery that the DB three-dimensional fold favors pseudoknot formation has implications for this element′s role in viral infection. In MBFVs, the DB is directly adjacent to the 3′ CS, a sequence responsible for the hybridization of 5′ and 3′ RNA elements and the subsequent genome cyclization required for viral replication. In the majority of MBFVs, with the exception of a clade of insect-specific viruses, the DB pseudoknot physically overlaps with the 3′ CS itself (de Borba et al., 2019). This suggests that DB pseudoknot formation is mutually exclusive with 5′-3′ CS base pairing, as has been demonstrated in SHAPE probing of a construct containing the DENV 5′ and 3′ UTRs (Sztuba-Solinska et al., 2013). The stability of the DB pseudoknot suggests that under the right conditions, the DB could act as an efficient competitor to 5′-3′ CS interactions, delaying replication timing until sufficient protein has been translated. Alternatively, a stable DB pseudoknot could also be a sensor of 5′-3′ cyclization, with cyclization inducing conformational changes within the DB and exposing DB sequences for potential RNA-protein interactions associated with replication.

Under the models presented here, the two DB pseudoknot conformations are connected with translation (pseudoknot formed) or genome replication (pseudoknot unformed). Given the high RNA sequence conservation within the DBs, it is tempting to speculate that protein co-factors could help regulate this equilibrium. Indeed, if the DB acts as a switch, RNA binding proteins could selectively stabilize or destabilize the pseudoknot to modulate this switch, with RNA binding helicases being an intriguing class of such proteins. In fact, such helicases have been previously implicated in DB binding and viral replication.

Specifically, host factor DDX6 in mammalian cells (Ward et al., 2011) and its insect homolog Me31B in mosquito cells (Goertz et al., 2019) are candidate RNA helicases that could promote viral replication by capturing the unfolded form or helping to unwind the DB pseudoknot. However, currently there is no direct evidence linking these helicases to changes in DB structure, and DDX6 and Me31B appear to perform distinct roles to either promote or restrict replication in mosquito vs mammalian cells (Goertz et al., 2019; Ward et al., 2011). Another candidate helicase is NS3, a virally produced protein involved in replication (Mazeaud et al., 2018). Overall, more research is needed to determine which factors may affect the structure of the DB during infection and how they relate to DB folding dynamics. Such insight will be useful to find ways to trap the DB in a single conformation, inhibiting a critical part of the viral life cycle.

In addition to the pseudoknot, the DBs contain a hyper-conserved region known as CS2. In our model, CS2 forms two helices (S3 and P3) within the four-way junction, likely imparting stability to the overall global RNA architecture. These helices are located adjacent to one another on a single side of the three-dimensional fold, forming a potential platform for interactions with proteins in the viral replication machinery that could explain the role of CS2 in promoting viral replication (Manzano et al., 2011). This RNA region was generally more reactive to the chemical probes, as both DENV-2 and DONGV DBs exhibited partial reactivity in their S3 regions, and this part of the RNA was involved in the intermolecular domain-swap in the crystal. Together, these observations suggest that CS2 may be inherently conformationally flexible. It is tempting to hypothesize that this flexibility may be conducive to protein binding and subsequent larger structural rearrangements. Given the connection of CS2 to the four-way junction and its potential to affect pseudoknot stability, binding of protein co-factors to CS2 could destabilize the overall architecture, connecting protein binding to pseudoknot unfolding and viral genome cyclization.

The 3′ UTRs of flaviviruses accumulate during infection as separate fragments known as sfRNAs, linked to cytopathic effects in infected cells and formed due to resistance of the 3′ UTR to 5′→3′ degradation by the host exonuclease XRN1 (Pijlman et al., 2008). As part of the 3′ UTR, DBs are contained both within the genomic RNA and in sfRNAs although we found no evidence that they block exonucleases to form sfRNAs. Currently it is unclear if the DBs perform a function as part of sfRNAs, however the fact that they are within the protected sfRNAs suggests they may have some other function in addition to their role in the viral genomic RNA.

## MATERIALS AND METHODS

### *In vitro* transcription

DNA templates for transcription were ordered as gBlocks from Integrated DNA technologies (IDT) and cloned into pUC19 and sequenced. DNA was amplified for transcription by PCR using custom DNA primers and Phusion Hot Start polymerase (New England BioLabs). Transcription reactions were conducted using 500 μL PCR reactions as template into 5 mL transcriptions. Transcription reactions contained: ∼0.1 μM template DNA, 8 mM each NTP, 60 mM MgCl_2_, 30 mM Tris pH 8.0, 10 mM DTT, 0.1% spermidine, 0.1% Triton X-100, and T7 RNA polymerase as well as 5 μL RNasin RNase inhibitor (Promega). Transcription reactions were left overnight, inorganic pyrophosphates were removed by centrifugation, and the reactions were ethanol precipitated and purified by denaturing PAGE purification. RNAs were eluted overnight at 4°C into ∼40 mL of diethylpyrocarbonate (DEPC)-treated milli-Q filtered water (Millipore) and concentrated using Amicon spin concentrators (Millipore).

### RNA crystallization and X-ray diffraction data collection

RNAs for crystallization were prepared as described above. The sequence used for *in vitro* transcription was 5′ - GGGGGCCAGATGTCATGTCTCTCAAGCCTAGGAGACACTAGACACTC TGGACTATCGGTTAGAGGAAACCCCCCCAAAAATGTATAGGCT<u>GAGGTGCTTGTATATAACCTCCACGA TGGTGCACCTTGGGCAACACTTCGGTGGCAAATCATCTACA</u> - 3′ where the underlined sequence represents a slightly modified sequence derived from the *Chilo* iridescent virus ribozyme (Webb and Luptak, 2011) which was used to generate homogenous 3′ ends. Ribozyme cleavage was facilitated at the end of the transcription reaction by adding MgCl_2_ (final conc. 120 mM) and incubating for 15 min at 65°C. The ribozyme-cleaved RNA was purified on a 10% dPAGE gel and re-folded at 65°C for 3 minutes at 5 mg/mL in a buffer containing 2.5 mM MgCl_2_ and 10 mM HEPES-KOH pH 7.5 and then allowed to equilibrate to room temperature before a final concentration of 0.5 mM spermidine was added. Crystal Screens I and II, Natrix I and II (Hampton Research) as well as the HELIX and MIDASplus screens (Molecular Dimensions) were used to perform initial screens at 20°C using single-drop vapor diffusion, and initial hits were optimized using custom screens. Crystals were grown by sitting drop vapor diffusion at 20°C in a solution containing in 50 mM Bis-Tris pH 7.0, 25 mM NaCl, 25 mM LiCl, 4.5 mM CaCl_2_, 50 mM Guanidine HCl, 23.75% polyethylene glycol 2000. Successful crystals appeared over several weeks. For data collection, crystals were gradually equilibrated into a freezing solution (crystallization solution supplemented to 27.5% polyethylene glycol 2000 + 4 mM Iridium(III) hexamine) and were flash-frozen using liquid nitrogen.

Diffraction data were collected at Advanced Light Source Beamline 4.2.2 using ′shutterless′ collection at the Iridium L-III edge (1.10208 Å) at 100°K. 180° datasets were collected with 0.2° oscillation images. Data were indexed, integrated, and scaled using XDS (Kabsch, 2010). SHELX (Schneider and Sheldrick, 2002) was used to obtain phasing information by single-wavelength anomalous dispersion (CFOM = 53.4).

### Model building and refinement

Iterative rounds of model building and refinement (simulated annealing, Rigid-body, B-factor refinement, occupancy) using COOT (Emsley and Cowtan, 2004) and Phenix (Adams et al., 2010) were used to generate a complete model of the RNA. Crystal diffraction data, phasing, and refinement statistics are contained in Table S1.

### RNA chemical probing

DNA templates for T7 RNA transcription follow. In all, <u>underlined</u> = T7 promoter, **bold** = normalization hairpins*, italics* = handle for FAM-labeled primer annealing

#### WT DONGV DB

<u>TAATACGACTCACTATA</u>**GGAAACGACTCGAGTAGAGTCGAAAA**TTAGGGGGCCAGATGTCATGTCTCT CAAGCCTAGGAGACACTAGACACTCTGGACTATCGGTTAGAGGAAACCCCCCCAAAAATGTATAGGCT ATTATATCGACA**GTTGGAGTCGAGTAGACTCCAACA***AAAGAAACAACAACAACAAC*

#### PKmut1 DONGV DB

<u>TAATACGACTCACTATA</u>**GGAAACGACTCGAGTAGAGTCGAAAA**TTAGGGGGCCAGATGTCATGTCTCT CAACGGATGGAGACACTAGACACTCTGGACTATCGGTTAGAGGAAACCCCCCCAAAAATGTATAGGCT ATTATATCGACA**GTTGGAGTCGAGTAGACTCCAACA***AAAGAAACAACAACAACAAC*

#### Pkmut2 DONGV DB

<u>TAATACGACTCACTATA</u>**GGAAACGACTCGAGTAGAGTCGAAAA**TTAGGGGGCCAGATGTCATGTCTCT CAAGCCTAGGAGACACTAGACACTCTGGACTATCGGTTAGAGGAAACCCCCCCAAAAATGTAATCCGT ATTATATCGACA**GTTGGAGTCGAGTAGACTCCAACA***AAAGAAACAACAACAACAAC*

#### PKcomp DONGV DB

<u>TAATACGACTCACTATA</u>**GGAAACGACTCGAGTAGAGTCGAAAA**TTAGGGGGCCAGATGTCATGTCTCT CAACGGATGGAGACACTAGACACTCTGGACTATCGGTTAGAGGAAACCCCCCCAAAAATGTAATCCGT ATTATATCGACA**GTTGGAGTCGAGTAGACTCCAACA***AAAGAAACAACAACAACAAC*

#### DENV-2 DB

<u>TAATACGACTCACTATA</u>**GGAAACGACTCGAGTAGAGTCGAAAA**CAACGGGGGCCAAGGTGAGATGAA GCTGTAATCTCACTGGAAGGACTAGAGGTTAGAGGAGACCCCCCGAAAACAAAAACAGCATATTGACG **GTTGGAGTCGAGTAGACTCCAACA***AAAGAAACAACAACAACAAC*

DNA templates were used for T7 RNA polymerase transcription as described above. Purified RNAs were subjected to chemical mapping procedures as previously described (Cordero et al., 2014). Briefly, 1.2 pmoles of RNA was re-folded and equilibrated to room temperature before adding chemical agents. In separate reactions RNA was probed using 5 μL of dimethyl sulfoxide (DMSO), 3 mg/mL N-methylisatoic anhydride (NMIA) or 1% dimethyl sulfate (DMS), and incubated at room temperature for 15–30 min.

Reactions were quenched using either 2-mercaptoethanol or 2-(N-morpholino) ethanesulfonic acid sodium salt (MES) pH 6.0. Chemically modified RNAs were isolated using a Poly(A)Purist™ MAG Kit (Thermo) and reverse transcribed using SuperScript III reverse transcriptase (Thermo) at 42 °C for 60 min using a fluorescently labeled primer (IDT): 5′ - /5-6FAM/ AAAAAAAAAAAAAAAAAAAAGTTGTTGTTGTTGTTTCTTT - 3′. Labeled DNA products were eluted in HiDi formamide spiked with Gene Scan ROX 500 size standard (Thermo). Samples were run on an Applied Biosystems 3500 XL capillary electrophoresis system and the data were analyzed using HiTRACE (https://ribokit.github.io/HiTRACE/) (Kladwang et al., 2011) with MatLab (MathWorks). HiTRACE measures the chemical reactivity at each nucleotide by reverse transcriptase stops. The reactivity due to treatment with DMSO was subtracted from the reactivity due to NMIA or DMS to subtract background signal.

Secondary structure diagram coloring was generated using MatLab and HiTRACE RiboKit: RiboPaint (https://ribokit.github.io/RiboPaint/tutorial/).

### Single-molecule FRET construct generation

Fluorescently labeled RNAs for smFRET were prepared as described (Akiyama and Stone, 2009). In brief, a Cy3-labeled RNA fragment was ligated to unlabeled 5′ and 3′ RNA fragments. The modified Cy3 labeled fragment was generated by labeling an amine-modified RNA containing a 5′ monophosphate ordered from Horizon Discovery with monoreactive Cy3 (GE Life Sciences) and the labeled RNA was purified by reverse-phase HPLC. The unlabeled 5′ fragment was ordered from Horizon Discovery. The unlabeled 3′ fragment was transcribed using T7 RNA polymerase and purified by denaturing polyacrylamide gel electrophoresis as described above. The 3′ fragment was then treated with recombinant BdRppH for 2 hours at 37°C in a buffer containing 50 mM Tris-HCl pH 7.5, 100 mM NaCl, 10 mM MgCl_2_, 1 mM DTT followed by Phenol:Chloroform:Isoamyl alcohol extraction and ethanol precipitation to convert the 5′-triphosphorylated RNA to 5′-monophosphorylated RNA (to facilitate ligation by T4 DNA ligase). The fluorescently labeled fragment was combined with the 5′ and 3′ unmodified RNAs by DNA-splinted RNA ligation using T4 DNA ligase (New England Biolabs) and purified by denaturing polyacrylamide gel electrophoresis. Fragments for splinted ligation are as follows:

5′ <u>fragment:</u> P – GGGGGCCAGAUGUC

<u>Middle fragment</u>: P – AUGUC[5-N-U]*CUCAA (* 5-aminoallyl-uridine)

3′ fragment Wild-type:

GCCTAGGAGACACTAGACACTCTGGACTATCGGTTAGAGGAAACCCCCCCAAAAATGTATAGGCTATTATATCGACACCTAACCACCAAGCCGACCG

3′ fragment PK mutant:

GCCTAGGAGACACTAGACACTCTGGACTATCGGTTAGAGGAAACCCCCCCAAAAATGTAATCCGTATTA TATCGACACCTAACCACCAAGCCGACCG

<u>DNA Splint</u>: AGTGTCTAGTGTCTCCTAGGCTTGAGAGACATGACATCTGGCCCCC

A biotin-labeled DNA handle (Integrated DNA Technologies) was ordered containing a 5′ biotin: 5′ - biotin - TTCGGTCGGCTTGGTGGTTAGGTGTCGA(iAmMC6T)**AT - 3′ (** Internal amine modified C6 dT) and was designed to hybridize with the RNA for surface immobilization and was labeled with Cy5 for FRET measurements. Cy5 labeling was accomplished by incubation with monoreactive Cy5 (GE Lifesciences) and DNA was purified by reverse-phase HPLC.

### smFRET data collection and analysis

Cy3-labeled purified RNAs were annealed to biotinylated Cy5-labeled DNA oligos and immobilized on PEGylated cover slides (Ted Pella, Inc.). PEGylated cover slides were prepared as described (Selvin and Ha, 2008). Slides were imaged on a total internal reflection fluorescence microscope (Nikon Eclipse Ti-E) using an Andor iXon+ DU897 CCD camera with a 100 msec integration time. The image was spectrally separated using the DV2 Dualview (Photometrics) and custom Semrock filters. FRET studies were performed in 100 mM NaCl, 50 mM Tris pH 7.5, 10% glycerol, 0.1 mg/mL BSA, 2 mM Trolox, 1% glucose, 1 mg/mL catalase, 1 mg/mL glucose oxidase and either 2 or 10 mM MgCl_2_. Analysis was performed using custom MATLAB (MathWorks) scripts.

### Structure-based sequence alignments

An initial list of 29 flaviviral DB sequences was manually compiled and a structural alignment was performed automatically using Locarna (https://rna.informatik.uni-freiburg.de/LocARNA/Input.jsp) and exported in Stockholm format. A search for structurally similar RNA sequences was performed using Infernal v. 1.1.2 (Nawrocki and Eddy, 2013) searching a database of *Flaviviridae* genomes from the NCBI viral genome browser (https://www.ncbi.nlm.nih.gov/genomes/GenomesGroup.cgi?taxid=11050). The consensus sequence and secondary structure model as well as the statistical analysis of covariation were calculated using the RNA Significant Covariation Above Phylogenetic Expectation (R-scape) program (Rivas et al., 2017) visualized using R2R (Weinberg and Breaker, 2011) and labeled in Adobe Illustrator.

Accession numbers for sequences used are as follows: NC_015843.2 Tembusu virus, NC_034151.1 T′Ho virus, NC_001477.1 Dengue virus 1, NC_006551.1 Usutu virus, NC_001563.2 West Nile virus lineage 2, NC_001474.2 Dengue virus 2, NC_018705.3 Ntaya virus, NC_000943.1 Murray Valley encephalitis virus, NC_001437.1 Japanese encephalitis virus, NC_012533.1 Kedougou virus, NC_009942.1 West Nile virus lineage 1, NC_002640.1 Dengue virus 4, NC_009028.2 Ilheus virus, NC_032088.1 New Mapoon virus, NC_009029.2 Kokobera virus, NC_007580.2 St. Louis encephalitis virus, NC_012532.1 Zika virus African isolate, NC_001475.2 Dengue virus 3,NC_012534.1 Bagaza virus, NC_035889.1 Zika virus Brazilian isolate, NC_009026.2 Bussuquara virus, NC_024017.1 Nhumirim virus, NC_002640.1 Dengue virus 4, NC_018705.3 Ntaya virus, NC_003635.1 Modoc virus, NC_034151.1 T′Ho virus, NC_001563.2 West Nile virus lineage 2, NC_009942.1 West Nile virus lineage 1, NC_006551.1 Usutu virus, NC_001475.2 Dengue virus 3, NC_007580.2 St. Louis encephalitis virus, NC_001477.1 Dengue virus 1, NC_015843.2 Tembusu virus flavivirus, NC_008718.1 Entebbe bat virus, NC_004119.1 Montana myotis leukoencephalitis virus, NC_009028.2 Ilheus virus, NC_009026.2 Bussuquara virus, NC_005039.1 Yokose virus, NC_033715.1 Nounane virus, NC_000943.1 Murray Valley encephalitis virus, NC_012735.1 Wesselsbron virus, NC_008719.1 Sepik virus, NC_017086.1 Chaoyang virus, NC_016997.1 Donggang virus, NC_002031.1 Yellow fever virus, NC_032088.1 New Mapoon virus.

### Protein expression

6x-histidine tagged *Kleuveromyces lactis* Xrn1 and *Bdellovibrio bacteriovorous* RNA pyrophosphate hydrolase (RppH) were expressed in BL21 *E. coli* cells and purified using nickel affinity and size exclusion chromatography by previously described methods (Steckelberg et al., 2018).

### Xrn1 digests

1 μg total RNA was resuspended in a 20 μL solution containing 50 mM Tris pH 8.0, 100 mM NaCl, 1 mM DTT, 10% glycerol, and either 1 mM or 10 mM MgCl_2_. The RNA was re-folded at 65°C for 5 minutes. 1 μl *Bdellovibrio bacteriovorus* RppH at a concentration of 0.5 μg/μL was added to all reactions and 1 μl *Kleuveromyces lactis* Xrn1 was added to every other reaction. Reactions were incubated for 1 hour at 37°C and separated on a 10% denaturing PAGE gel and visualized by staining with ethidium bromide. DNA constructs for RNA transcription for Xrn1 digestion were as follows:

#### Zika virus Xrn1-resistant RNA + leader

GAAGCACCAATCTTAATGTTGTCAGGCCTGCTAGTCAGCCACAGCTTGGGGAAAGCTGTGCAGCCTGTGACCCCCCCAGGAGAAGCTGGGAAACCAAGCCTAT

#### Zika virus DB RNA + leader

GAAGCACCAATCTTAATGTTCAATCTGGGGCCTGAACTGGAGATCAGCTGTGGATCTCCAGAAGAGGGACTAGTGGTTAGAGGAGACCCCCCGGAAAACGCAAAACAGCATA

### Northern blots

#### Viral infection

A549 cells were plated into 6-well plates so that at the time of harvest each well would have approximately 1 million cells. For infection, 2 mL of culture media was aspirated from each well, followed by a wash with 1 mL of 1X PBS. Each well was infected with 500 μL of ZIKV viral inoculum at an MOI of 3. Plates were incubated with viral inoculum at 37°C for one hour. After incubation viral inoculum was removed and 2 mL of complete Ham′s F-12 media was added to each well. After 48 hours, cells were harvested. For harvest, media was aspirated from the plate. Each well was washed once with 1 mL 1X PBS and RNA was extracted using the EZNA Total RNA kit (Omega Biotek). 375 μL of TRK lysis buffer with β-Mercaptoethanol was added to each well and incubated for 3 minutes at room temperature. Two biological replicates were combined into one sample for RNA extraction (each RNA extraction sample was from an approximate total of 2 million cells) and the RNA was column purified via manufacturer′s instructions.

#### Preparation of probes

A 35-mer DNA oligo complementary to nucleotides 263-298 of the PRVABC59 strain 3′ UTR (sequence: 5′- TCCTCTAACCACTAGTCCCTCTTCTGGAGATCCAC -3′) as well as a DNA oligo complementary to nucleotides 323-358 (sequence: 5′ - CTCATGGAGTCTCTGGTCTTTCCCAGCGTCAATAT - 3′) and complementary to the U6 snRNA (sequence: 5′ - TATGGAACGCTTCACGAATTTGCGTGTCATCC - 3′) was ordered from IDT. 100 pmol of DNA was incubated with 2 μL 5mCi [γ32-P]ATP (Perkin Elmer) and 4 μL T4 polynucleotide kinase (New England BioLabs) in a 100 μL reaction for 2 hours and purified using P-30 spin columns (BioRad). The radiolabeled probe was heated to 100°C for 2 minutes and resuspended in 10 mL of ULTRAhyb Oligo hybridization buffer (Ambion). Blots were incubated with 10 mL of the resuspended probe overnight at 42°C.

#### Blotting procedure

1 μg of total RNA from mock- and ZIKV-infected cells was resuspended in 2x formamide RNA loading buffer and run on a 6% denaturing PAGE gel (Invitrogen) with an RNA ladder (Thermo Scientific). Gels were stained with ethidium bromide and imaged to obtain the position of the RNA ladder. The RNA from the gels was subsequently transferred to a HyBond-N+ nylon membrane (GE Life Sciences) using an electrophoretic transfer apparatus (Idea Scientific). The membrane was crosslinked using a UV stratalinker and blocked at 42°C using ULTRAhyb Oligo hybridization and blocking buffer (Ambion) for 2 hours while rotating. Blots were probed rotating at 42°C with a 35-mer DNA oligo prepared as described above overnight and washed in 2x saline-sodium citrate (SSC) buffer with 0.5% SDS for 10 minutes at 42°C four times. The blots were imaged with a phosphor screen and a Typhoon scanner (GE Life Sciences) and were aligned with the ethidium-stained image of the gels to obtain the position of size standards from the RNA ladder.

## DATA DEPOSITION

The coordinates have been deposited in the Protein Data Bank with the accession code XXXX (to be supplied prior to publication).

## ACKNOWLEDGEMENTS

We thank the members of the Kieft Lab for useful discussions and insight, and Anna-Lena Steckelberg, Steve Bonilla Rosales, and David Costantino for critical reading of the manuscript. This work was funded by grants NIH R35GM118070 to JSK, and Department of Defense MIIRA PRMRP PR160117 and VA Merit Award I01BX003863 to JDB. We thank Jay Nix (ALS Beamline 4.2.2) and John Hardin (UC Anschutz Medical Campus) for their assistance with data collection. Beamline 4.2.2 at the Advanced Light Source is partially funded by the NIH (P30GM124169-01) and operated under contract with the U.S. DOE(DE-AC02-05CH11231). The University of Colorado Anschutz Medical Campus x-ray crystallography facility is supported by the NIH (P30CA046934 and S10OD012033).

---

**Figure 1 – figure supplement 1.**
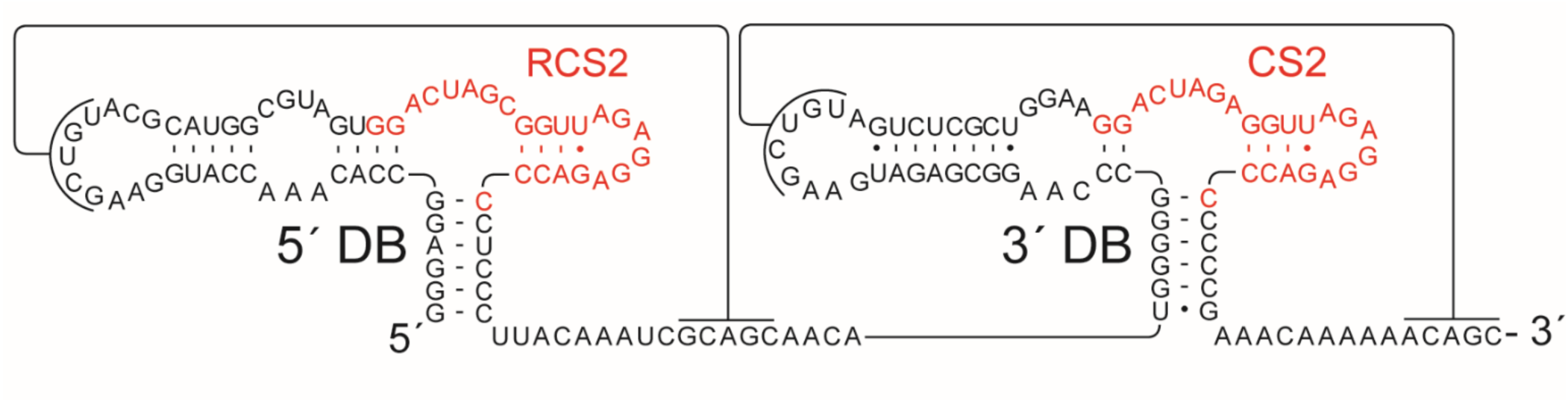
Organization of RCS2 and CS2 motifs. Secondary structure diagram of DENV-2 (accession number NC_001474.2) DB region showing the 5′ and 3′ DBs. The locations of the RCS2 and CS2 motifs are highlighted in red.

**Figure 1 - figure supplement 2.**
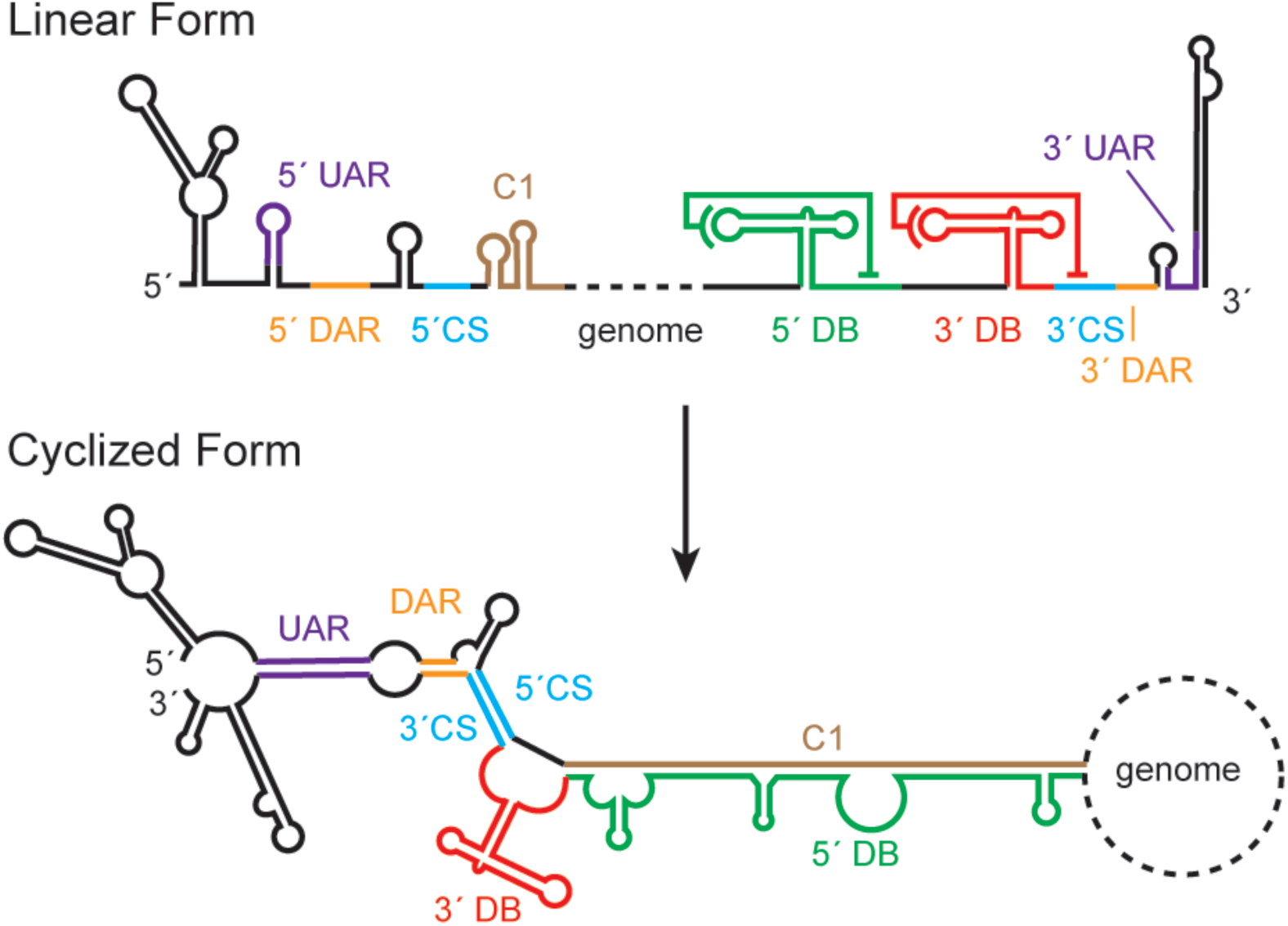
Proposed genome rearrangements during viral cyclization. The 3′ DB (red) pseudoknot unfolds during viral cyclization, exposing the 3′ CS for binding to the 5′ CS (cyan). 5′-3′ CS base pairing is supplemented by base pairing in the downstream of AUG region (DAR, orange) and upstream of AUG region (UAR, magenta). The 5′ DB (green) partially unfolds to base pair with the C1 region (brown). Figure derived from (de Borba et al., 2015).

**Figure 1 – figure supplement 3.**
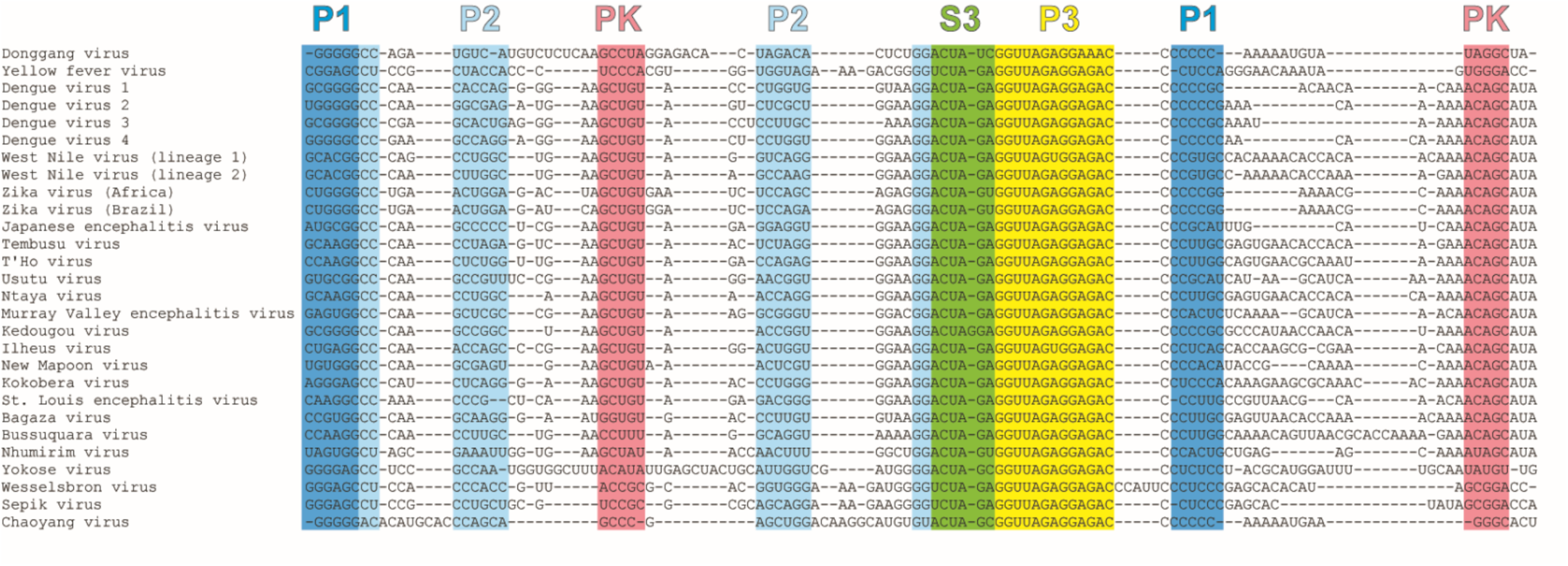
Structural conservation of DONGV to other MBFVs. Sequence alignment of MBFV DB RNAs demonstrating sequence homology of DONGV to related MBFVs. Structure elements from the DONGV crystal structure are highlighted in the alignment and colored as in Figure 2.

**Figure 2 – figure supplement 1.**
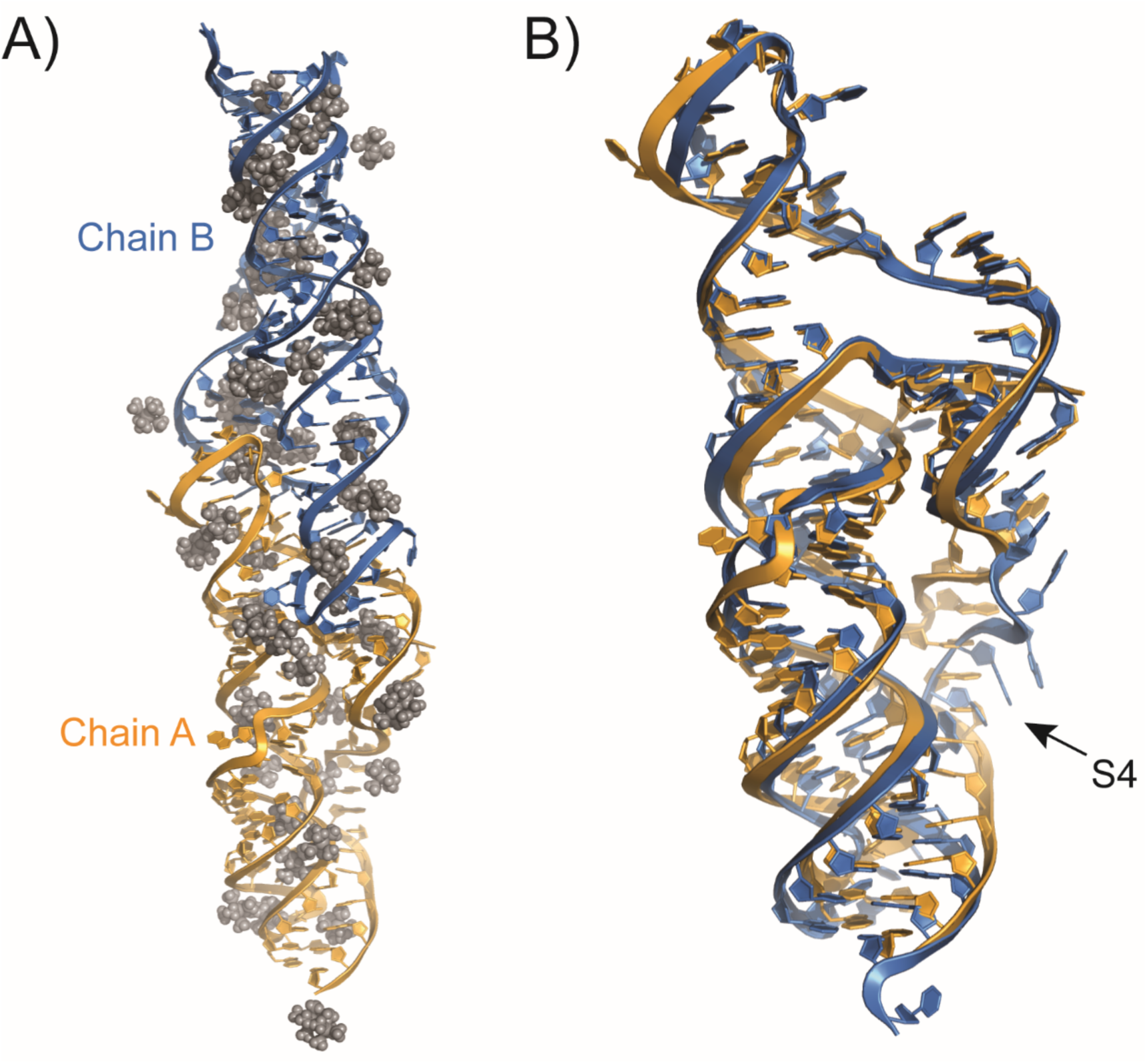
Comparison of two molecules in the asymmetric unit. **A)** The final refined structure of the crystallographic asymmetric unit containing two copies of the RNA (blue and orange) with positions of iridium (III) hexammine shown in grey. **B)** Both copies of the RNA in the asymmetric unit are shown overlaid, with an overall RMSD of 2.3 Å (all atoms).

**Figure 2 – figure supplement 2.**
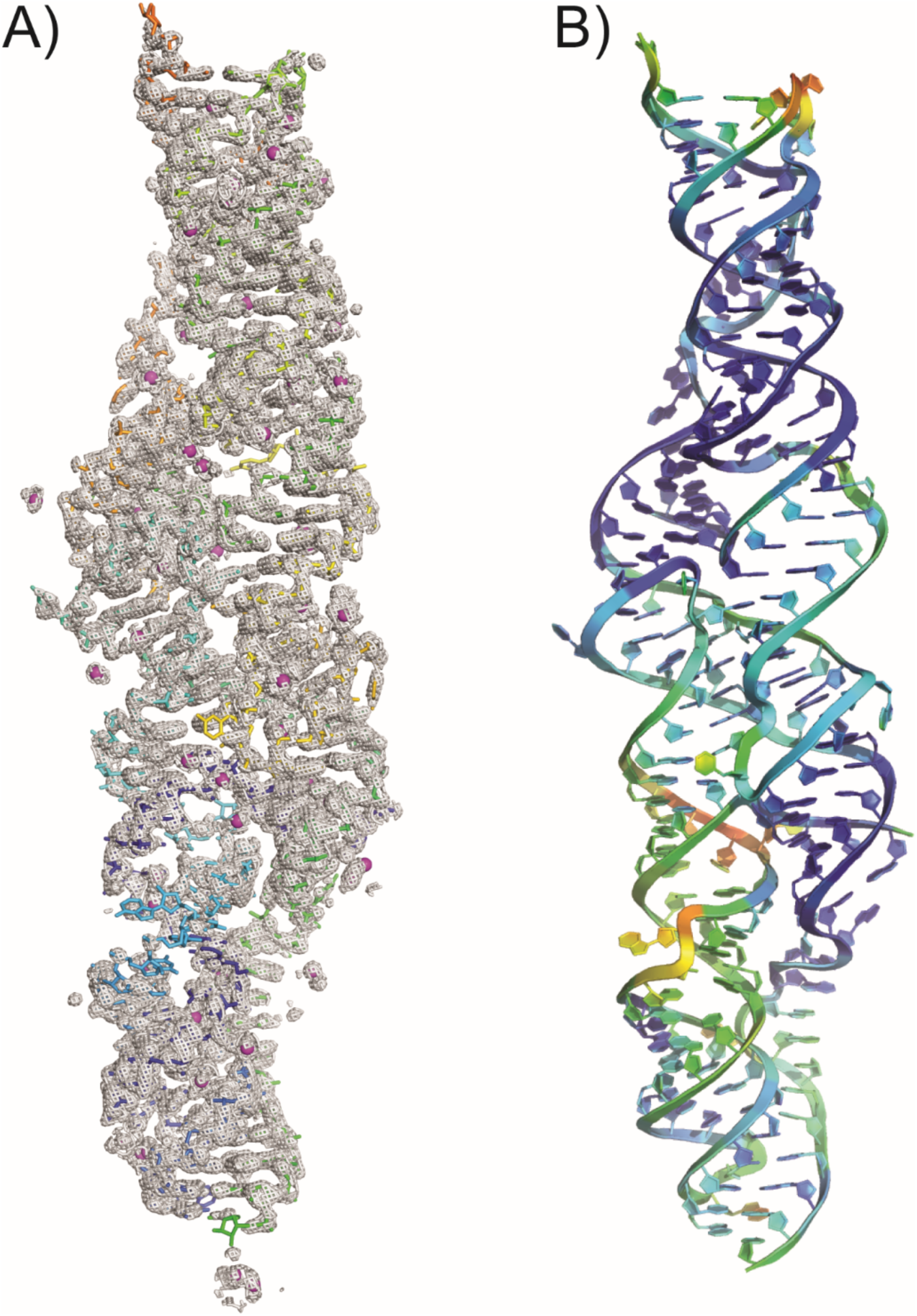
Composite omit map and relative b-factors. **A)** Simulated annealing composite omit map of the asymmetric unit of the DONGV DB RNA crystal structure at a 1 σ cutoff compared with the three-dimensional structure model. Iridium atoms are shown as magenta spheres with amines omitted for clarity. **B)** The two copies of the RNA in the asymmetric unit colored by relative B-factors, red represents highest and blue represents lowest.

**Figure 4 – figure supplement 1.**
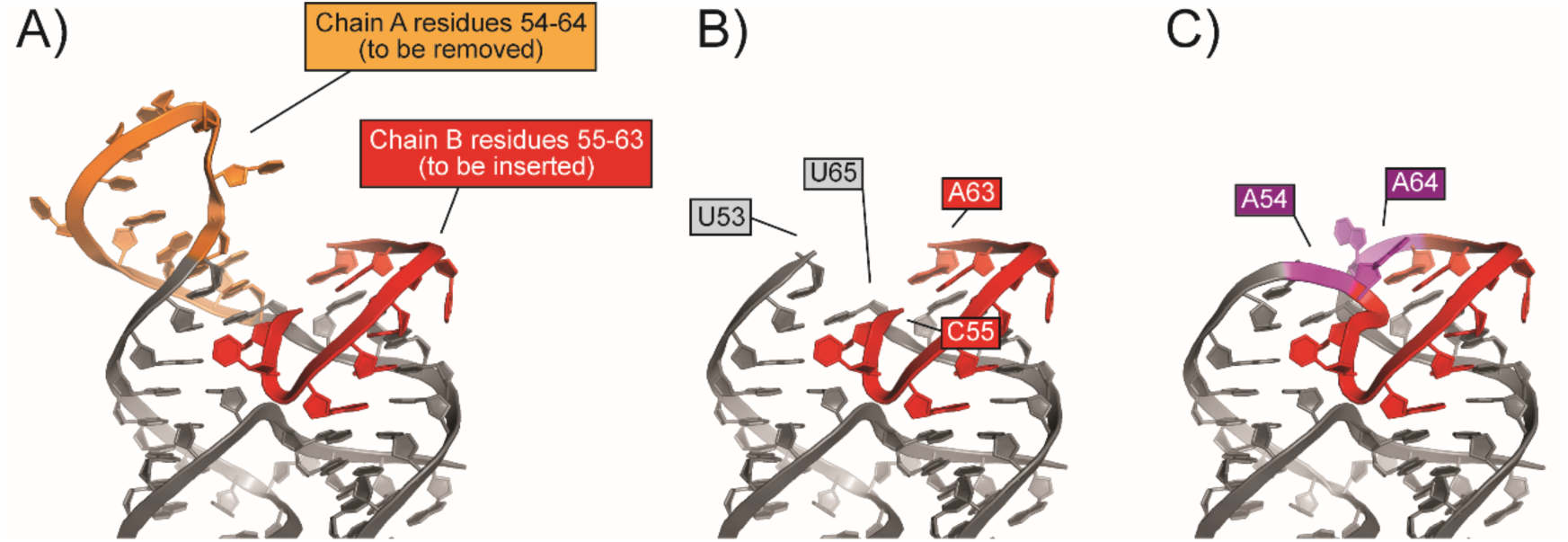
Modeling of the monomeric solution structure. **A)** Given the observed base pairing between residues 57-60 and 67-70 *in trans*, we determined residues 55-63 from chain A (orange) could be replaced with residues 55-63 from Chain B (red) to obtain a model of a solution structure of the DONGV DB RNA. **B)** Model of the monomeric structure of the DONGV DB lacking connecting residues A54 and A64. **C)** To complete the model, residues A54 and A64 (magenta) were inserted and energy minimized using Coot without electron density restraints in order to bridge U53 and U55 as well as G65 and A63 respectively.

**Figure 5 – figure supplement 1.**
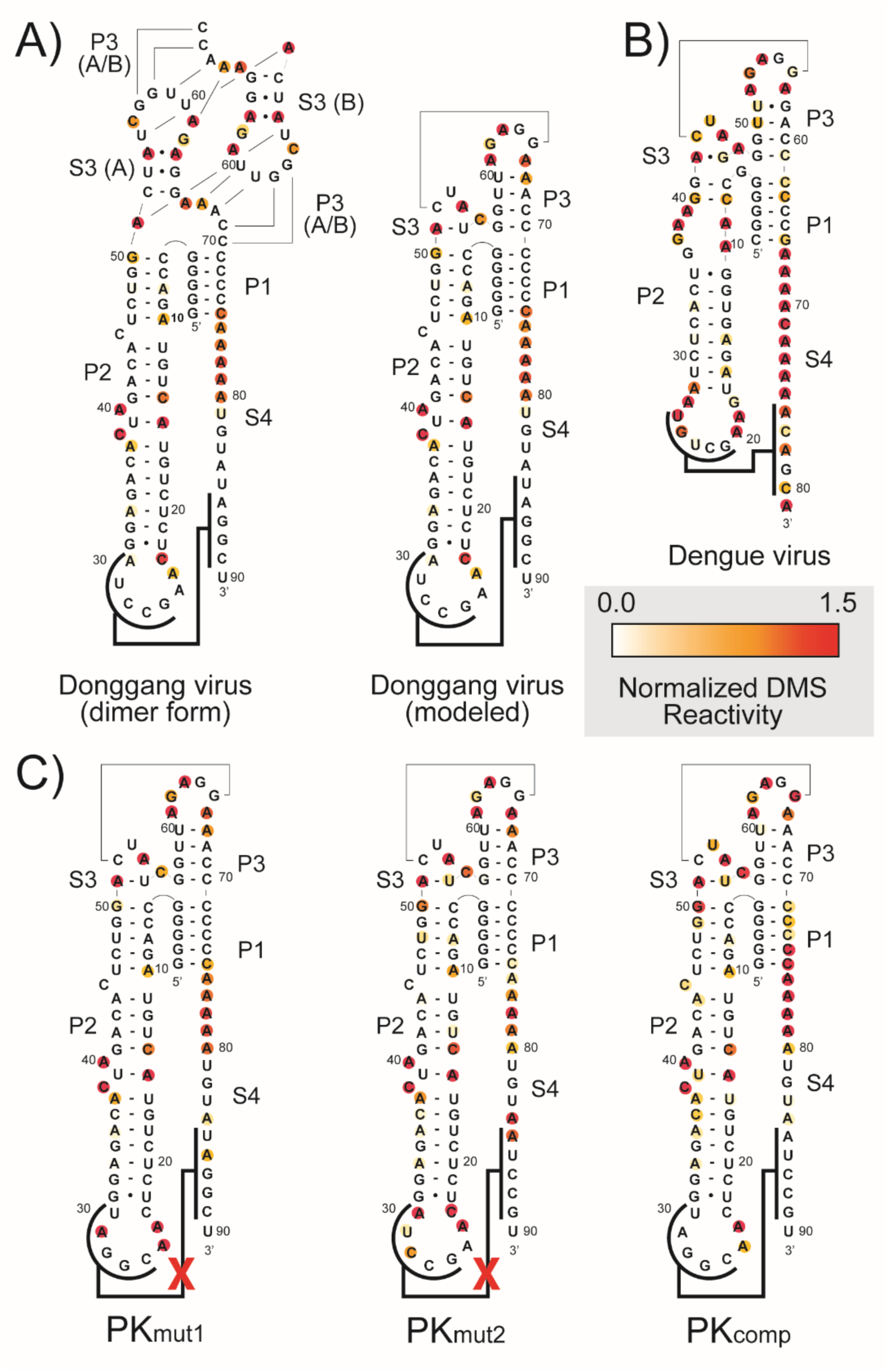
DMS chemical probing of the DB RNA. Normalized DMS reactivity plotted against the secondary structure of A) DONGV DB crystal and modeled forms, B) DENV-2 DB, and C) DONGV pseudoknot mutants.

**Figure 5 – figure supplement 2.**
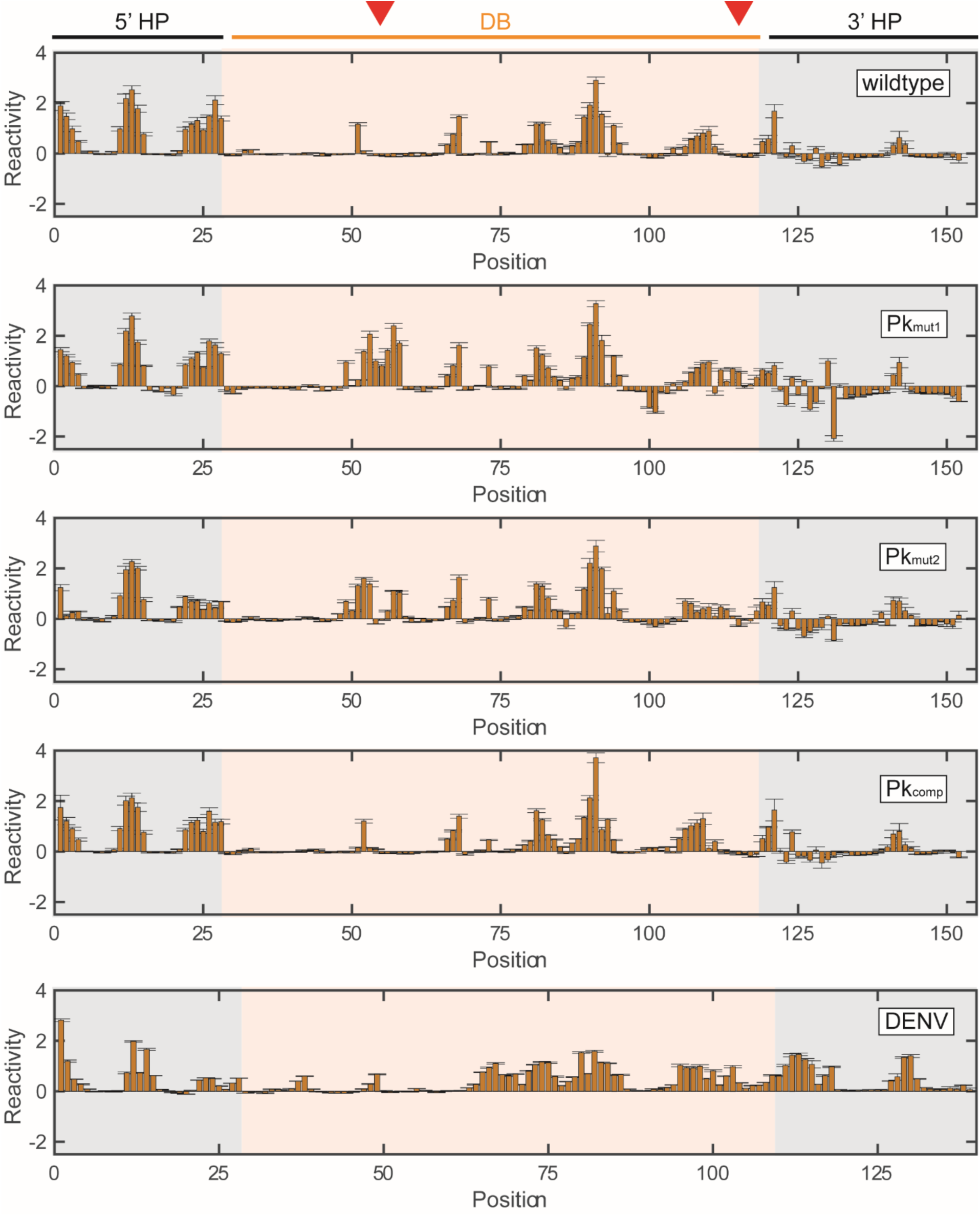
Quantitation of NMIA chemical probing experiments. NMIA chemical probing for DONGV wildtype, PK_mut1_, PK_mut2_, and PK_comp_ DB constructs as well as the DENV-2 DB. Bars represent the difference in average reactivity from 4 replicate experiments between NMIA-treated RNA and a DMSO-treated control. Error bars = standard deviation. The position of the DB RNA within the construct is indicated in orange, as well as the position of 5′ and 3′ hairpins (HP) used for normalization and to confirm proper RNA folding in grey. Red triangles indicate the position of the pseudoknot.

**Figure 5 – figure supplement 3.**
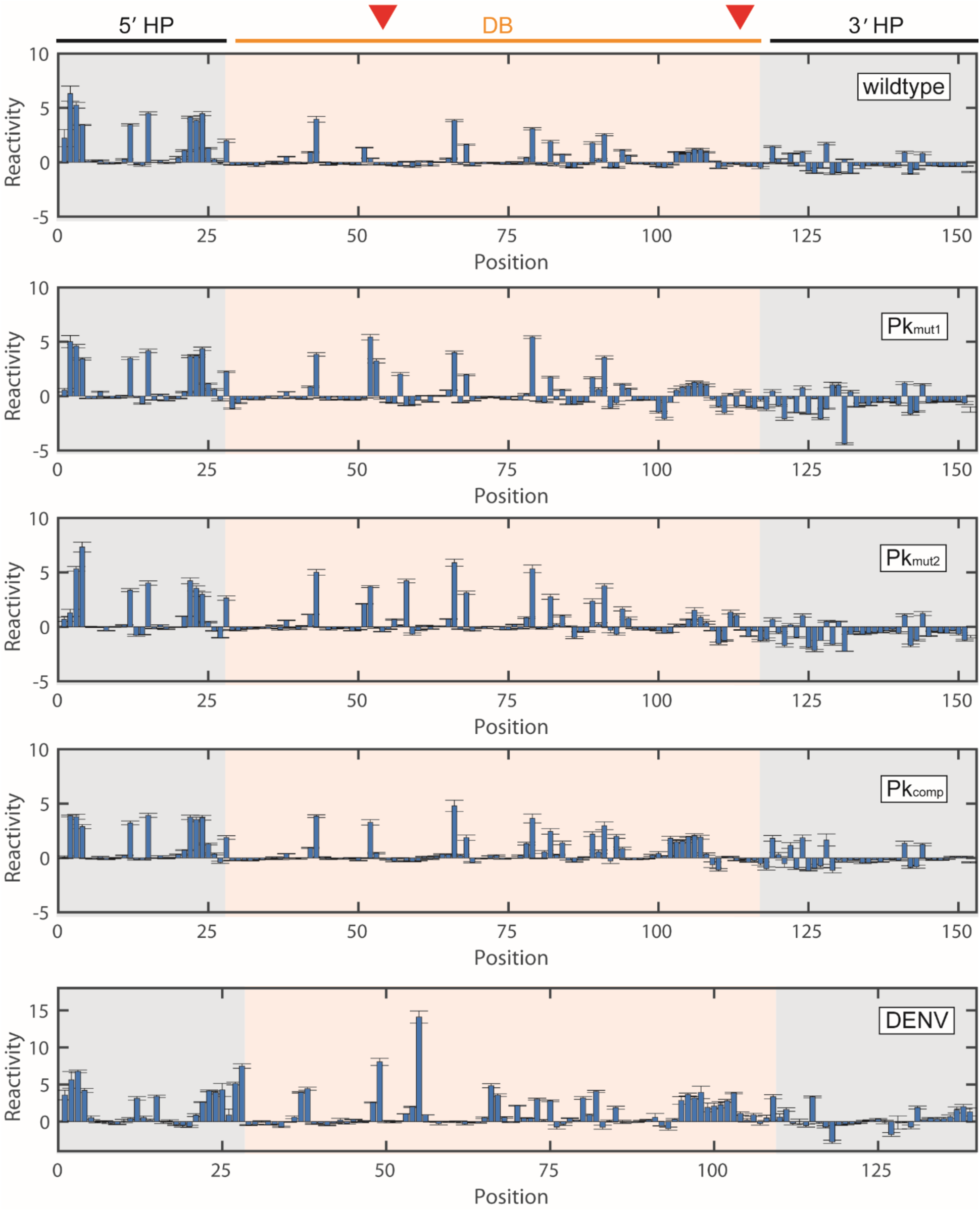
Quantitation of DMS chemical probing experiments. DMS chemical probing for Donggang wildtype, PK_mut1_, PK_mut2_, and PK_comp_ DB constructs as well as the DENV-2 DB. Bars represent the difference in average reactivity from 4 replicate experiments between DMS-treated RNA and a DMSO-treated control. Error bars = standard deviation. The position of the DB RNA within the construct is indicated in orange, as well as the position of 5′ and 3′ hairpins (HP) used for normalization and to confirm proper RNA folding in grey. Red triangles indicate the position of the pseudoknot.

**Figure 6 – figure supplement 1.**
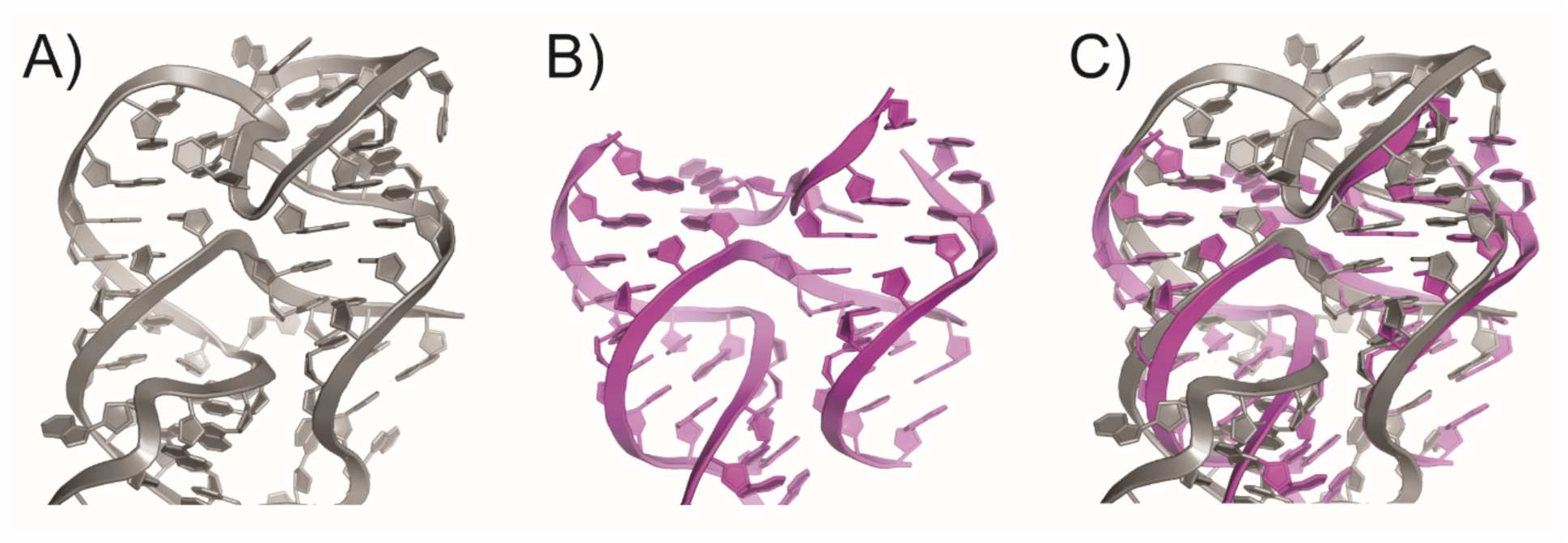
Structural comparison of homologous four-way junctions. **A)** Four-way junction of the DONGV DB RNA. **B)** Four-way junction from RNase P (PDB ID: 1NBS) **C)** Structural superposition of (A) and (B).

**Figure 7 – figure supplement 1.**
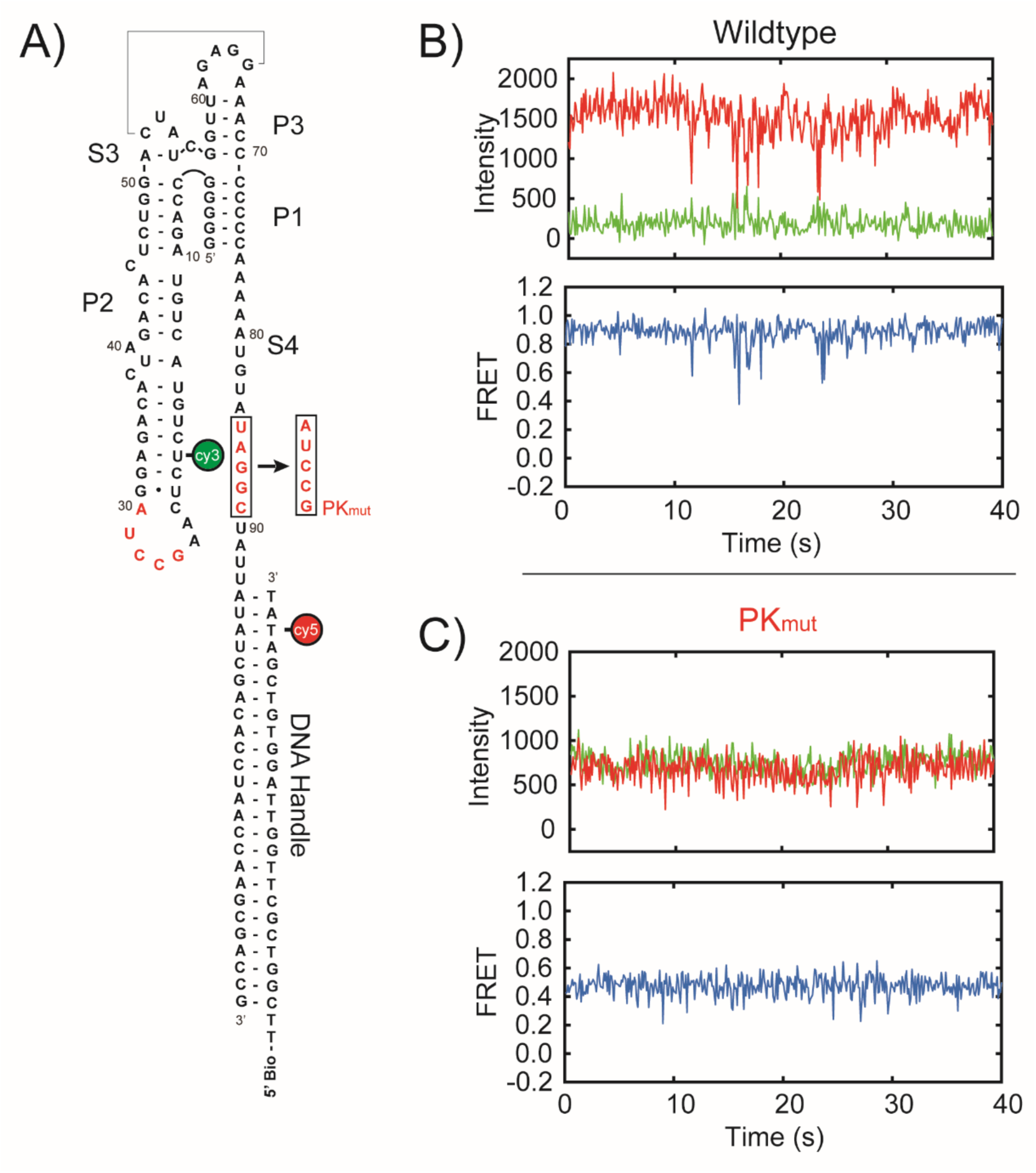
Overview of smFRET experiments. **A)** Schematic of complete sequences used in smFRET experiment. Cy3 was site-specifically incorporated on residue U20 of the DONGV DB sequence and Cy5 was incorporated on the DNA handle. The DNA handle was modified with a 5′ biotin in order to immobilize the complex on a microscope slide by a biotin-streptavidin linkage. Pseudoknot residues are highlighted in red and the sequence of the pseudoknot mutant (PK_mut_) is shown. **B)** Representative single molecule trace of a wildtype DONGV DB RNA. (Top) Intensity of Cy3 (green) and Cy5 (red) over time. (Bottom) Calculated FRET values over time. **C)** Representative single-molecule trace of the PK_mut_ construct showing Cy3 and Cy5 intensity and calculated FRET values as in B.

**Figure 7 – figure supplement 2.**
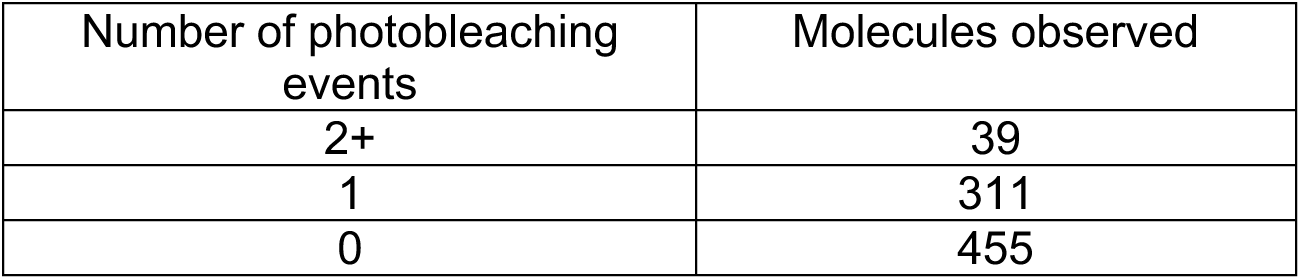
Tabulation of observed photobleaching events. Counting of double, single, and no photobleaching events observed in 160 second smFRET traces.

**Figure 7 – figure supplement 3.**
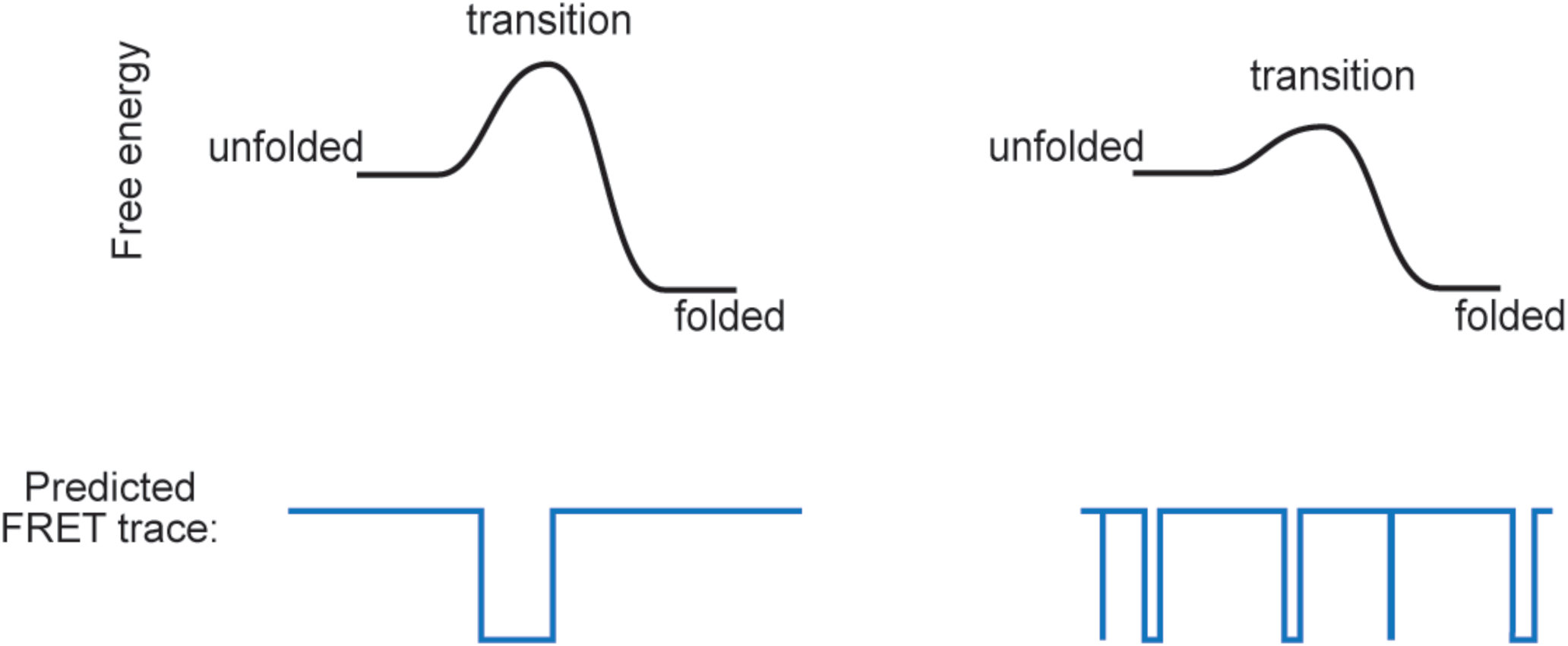
Free energy landscapes and predicted smFRET trace values. A molecule with a high free energy to the transition state(left) will have long dwell times in the folded and unfolded states. A molecule with a lower free energy to the transition state (right) will have more frequent FRET changes with shorter dwell times. The FRET behavior of the DONGV DB RNA more closely resembles the example of the low free energy transition state. Note that the free energy difference between the folded and unfolded states remains the same in both cases, with the folded state predominating.

**Figure 7 – figure supplement 4.**
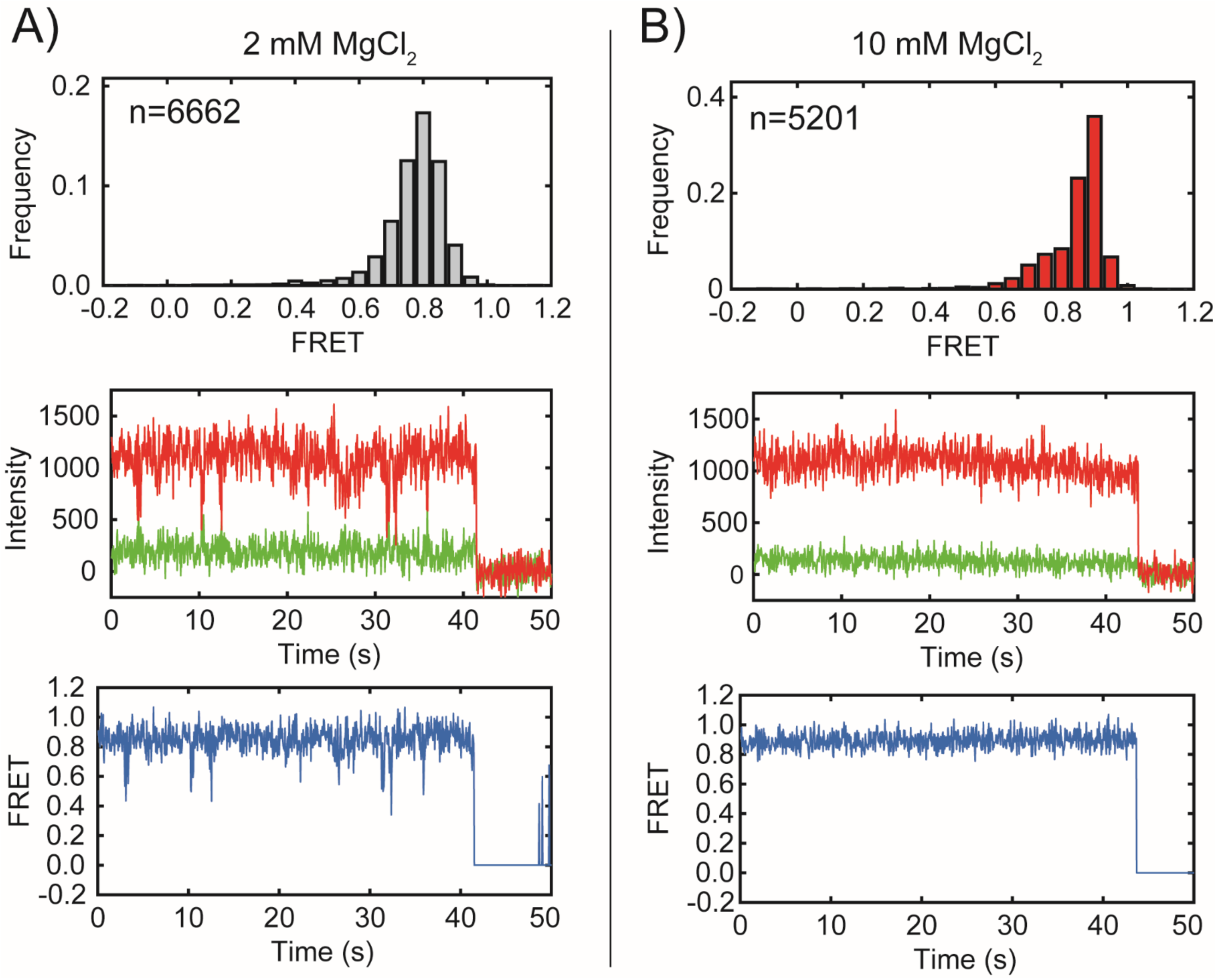
Magnesium dependence of pseudoknot stability. **A)** Single-molecule histogram (top) of the FRET distribution of 6662 wildtype DONGV DB RNAs with dyes placed as in (Figure 7) in 2 mM MgCl_2_. A representative single-molecule FRET trace following an individual molecule is below. The trace shows Cy3 (green) and Cy5 (red) intensity over time, with a calculated FRET trajectory (blue) displayed below. The loss of signal at the end of the trace is due to Cy3 photobleaching under laser illumination. **B)** Single-molecule histogram (top) of the FRET distribution of 5201 wildtype DONGV DB RNAs as in **(A)** but instead in 10 mM MgCl_2_. A representative single-molecule FRET trace following an individual molecule is below and matches the scheme shown at left.

